# Supervised dimension reduction for large-scale “omics” data with censored survival outcomes under possible non-proportional hazards

**DOI:** 10.1101/586529

**Authors:** Lauren Spirko-Burns, Karthik Devarajan

## Abstract

The past two decades have witnessed significant advances in high-throughput “omics” technologies such as genomics, proteomics, metabolomics, transcriptomics and radiomics. These technologies have enabled simultaneous measurement of the expression levels of tens of thousands of features from individual patient samples and have generated enormous amounts of data that require analysis and interpretation. One specific area of interest has been in studying the relationship between these features and patient outcomes, such as overall and recurrence-free survival, with the goal of developing a predictive “omics” profile. Large-scale studies often suffer from the presence of a large fraction of censored observations and potential time-varying effects of features, and methods for handling them have been lacking. In this paper, we propose supervised methods for feature selection and survival prediction that simultaneously deal with both issues. Our approach utilizes continuum power regression (CPR) - a framework that includes a variety of regression methods - in conjunction with the parametric or semi-parametric accelerated failure time (AFT) model. Both CPR and AFT fall within the linear models framework and, unlike black-box models, the proposed prognostic index has a simple yet useful interpretation. We demonstrate the utility of our methods using simulated and publicly available cancer genomics data.

## 1 Introduction

Advances in high-throughput technologies in the past two decades have enabled large-scale “omics” studies that generate enormous amounts of data that are measured on a variety of scales. Examples include, but are not limited to, genomic studies such as next-generation sequencing, methylation, allele-specific expression, microarrays, and DNA copy number as well as studies involving genome-wide association, proteomics, metabolomics, transcriptomics and radiomics. Genomic studies, for instance, enable the simultaneous measurement of the expression profiles of tens of thousands of genomic features, often from a relatively small number of individual patient samples. Such studies result in massive quantities of data requiring analysis and interpretation while offering tremendous potential for growth in our understanding of the pathophysiology of many diseases. When information on an outcome variable such as time to an event (or survival time) is available, one of the goals of an investigator is to understand how the expression levels of genomic features, and clinical and demographic variables (covariates) relate to an individual’s survival over the course of a disease. The number of covariates (*n*) far exceeds the number of observations (*p*), typically, in these large-scale genomic studies. With the tremendous volume of information available, the investigator can now estimate and attempt to understand the effects of specific genomic features on various diseases with the ultimate goal of developing a prognostic profile of patient survival. In this context, biomarker discovery poses many challenges and plays a pivotal role in the search for more precise treatments. The role and significance of the analysis of time-to-event data in cancer research cannot be overstated where current efforts focus on predicting therapeutic responses of patients with a view to personalizing cancer treatment.

The ill-conditioned problem of predicting the survival probability when *p* > > *n* is further compounded by the presence of censored survival times. In this high-dimensional setting, one is often interested in building a genomic profile that is predictive of the survival probability for a new patient. The Cox proportional hazards (PH) model is the most celebrated and widely used statistical model linking survival time to covariates (Cox, 1972). It is a multiplicative hazards model that implies constant hazard ratio (HR); that is, it postulates that the risk (or hazard) of death of an individual given their covariates is simply proportional to their baseline risk in the absence of any covariate. While this model has proved to be very useful in practice due to its simplicity and interpretability, the assumption of constant HR has been shown to be invalid in a variety of situations in medical studies (Devarajan & Ebrahimi, 2011; Peri et al., 2013). When applied to our problem, the PH model would implicitly assume a constant effect of genomic feature expression on survival over the entire period of follow-up in a study, a supposition that is neither verifiable nor likely for each feature. For example, non-proportional hazards (NPH) can occur when the effect of a genomic feature increases or decreases over time leading to converging or diverging hazards (CH or DH), a situation that cannot be handled by the PH model (Bhattacharjee et al., 2001; Xu et al., 2005; Dunkler et al., 2010; Rouam et al., 2011). In addition, NPH can result from model misspecification such as from omitting a strong clinical covariate (for instance, age at diagnosis or stage of disease) or another genomic feature. Another scenario encountered in practice is the case of dependence between covariates and the censoring time distribution (Chen et al., 2002). These scenarios require more general survival models that consider time-varying covariate effects. Examples of such models include the Accelerated Failure Time (AFT) and Proportional Odds (PO) models, among others (Buckley & James, 1979; Jin et al., 2006; Martinussen & Scheike, 2006; Yang & Prentice, 2005; Devarajan & Ebrahimi, 2011). The AFT model is a censored linear regression model in which the covariates cause an acceleration (linear transformation) of the time scale while the PO model postulates that the odds of death for an individual, given their covariates, is simply proportional to their baseline odds - a situation typically encountered when the effect of a genomic feature decreases with time leading to diverging hazards. Unlike PH, these models do not imply a constant HR and, interestingly, both PH and PO models intersect with the AFT model. Moreover, the AFT model can accommodate a variety of well known survival time distributions - such as the lognormal, log-logistic, Weibull and exponential, to name a few - useful for modeling censored survival data in practice or accommodate a completely distribution-free approach with no prior assumption on the data generating mechanism (Kalbfleisch & Prentice, 2002; Jin et al., 2006). These attractive properties provide modeling flexibility and make the AFT model a versatile alternative useful for handling a variety of data structures.

As evidenced by the following literature survey, very little research has been done to account for the time-varying effect of genomic features or to study the consequences of NPH on feature selection and survival prediction, despite its clear importance in translational medicine. Within a broader context, these shortcomings extend to the many types of high-throughput “omics” studies outlined earlier. In this paper, we generically use the term “omics” to represent this variety and the term feature to denote the appropriate “omic” feature of interest. The rest of the paper is organized as follows. In *§*2, we survey existing methods for supervised dimension reduction within the context of high-throughput “omics” data and censored survival outcomes, and discuss their weaknesses. Section 3 begins with a motivation of the need for a flexible method using real-life cancer genomic data. In *§*4, we propose an approach that combines continuum power regression with the parametric or semi-parametric AFT model and in *§*5, we develop a prognostic index and survival prediction algorithm using this approach. Section 6 is devoted to simulation studies for evaluating the proposed methods while *§*7 focuses on the application of these methods to several publicly available data sets in cancer genomics. Last but not least, *§*8 provides a summary and discussion including future work. The Supplementary Information (SI) section contains detailed results from simulation studies and real-life data sets.

## 2 A brief survey of existing methods and their limitations

A variety of methods are currently available in the literature for handling a large number of features in conjunction with censored survival outcomes. These include methods based on principal components regression (PCR) (Li & Li, 2004; Bair et al., 2006; Ma et al., 2006), partial least squares (PLS) (Park et al., 2002; Li & Gui, 2004; Nguyen & Rocke, 2002; Nguyen, 2005; Nygard et al., 2008; Boulesteix & Strimmer, 2007; Devarajan et al., 2010; Bastien et al., 2015) and regularization such as the ridge regression, least absolute shrinkage and selection operator (LASSO), least angle regression, elastic net or related variants (Tibshirani, 1997; Gui &, 2005; Segal, 2008; Engler & Yi, 2009; Wang et al., 2009; Kaneko et al., 2012). Other available methods in this context include those based on boosting (Li & Luan, 2005; Wei & Li, 2007; Luan & Li, 2008; Lu & Li, 2008), sure screening procedures (Fan et al., 2010), Cox assisted clustering (Eng & Hanlon, 2012), networks (Zhang et al., 2013), kernel methods (Li & Luan, 2003) and nested cross-validation (Laimighofer et al., 2016).

A comparison of various existing methods has revealed that those based on a linear combination of features (such as PCR and PLS) or regularization (such as LASSO etc.) showed overall superior performance (Bovelstad et al., 2007; van Wieringen et al., 2009; Witten & Tibshirani, 2008). Methods based on PLS and PCR typically utilize all features for prediction and cannot directly specify relevant features that are associated with survival. Regularization methods generally perform well in this setting by identifying unimportant features from the large number of features present by shrinking their coefficients to exactly zero. However, a method such as LASSO suffers from some fundamental limitations due to the *L*_1_ penalty. For instance, the number of non-zero coefficients can be at most *n*, i.e., the number of features that can be selected by LASSO is bounded by the sample size of the data set (Rosset et al., 2004). In large-scale genomic studies, this can lead to the unrealistic conclusion that no more than *n* genomic features are relevant to survival in a complex biological process where *p* ≫ *n* are actually present. This is further compounded by the relatively small number of observations often seen in these biomedical studies. Moreover, the expression levels of features sharing a particular biological pathway can be highly correlated. It is therefore desirable to have a method that automatically selects the entire set of correlated features; however, LASSO can usually select only one feature in this situation. On the other hand, a method such as ridge regression necessarily selects all features in a data set. These issues can pose serious problems particularly when dealing with the ultra high-dimensional data sets obtained in modern “omics” studies (Wang et al., 2008).

An inherent weakness of these methods is the assumption of PH in their formulation which does not permit the incorporation and, therefore, the detection of time-dependent covariate effects. However, there exist methods based on alternate survival models such as the AFT (Huang et al., 2006; Wang & Leng, 2007; Datta et al., 2007; Wang et al., 2008; Luan & Li, 2008; Cai et al., 2009; Wang & Wang, 2010; Engler & Yi, 2009; Liu et al., 2010; Devarajan et al., 2010), PO (Lu & Li, 2008), non-linear transformation (Lu & Li, 2008), additive hazards (Ma et al., 2006) or that are model-free (Van Belle et al., 2011; Geng et al., 2014; Pang et al., 2012). Huang et al. (2006) combine the AFT model with LASSO or threshold-gradient-directed regularization (TGDR) using Stute’s estimator (Stute, 1993), thereby providing flexible methods for handling NPH and high-dimensionality. In addition to known limitations, their LASSO approach has been known to result in inferior prediction accuracy in empirical studies. Furthermore, TGDR is sensitive to the choice of a parameter value that could significantly alter the number of features selected and thus lead to overestimation of the number of non-zero coefficients, potentially further reducing the number of features selected (Wang & Wang, 2010). Datta et al. (2007) developed an approach that combines standard PLS or LASSO with the AFT model after mean imputation of censored observations. This approach does not improve upon these existing methods and suffers from the limitations of LASSO. Devarajan et al. (2010) outlined a PLS-based method for lognormally distributed data; however, it not only relies on unrealistic assumptions on the data generating mechanism but also cannot be used on independent test data. Wang et al. (2008) proposed a doubly penalized method based on the AFT model for estimation, feature selection and survival prediction by extending elastic net regression for linear models to censored survival data. Unlike LASSO, this approach can select an arbitrary number of highly correlated features with non-zero coefficients; however, it involves the selection of tuning parameters and can be computationally slow. Engler & Yi (2009) proposed an elastic net approach with mean imputation in conjunction with the Cox PH or AFT model and showed that the AFT version showed better performance. Existing model-free methods provide a flexible alternative that can account for linear and non-linear covariate effects; however, they tend to be computationally infeasible and typically require the choice of various tuning parameters.

## 3 Motivation for the proposed methods

In the high-dimensional setting, incorporating time-dependent covariates in the Cox PH model, use of stratification or separate modeling for different time periods in order to account for NPH are prohibitive and infeasible. As outlined in the literature survey above, computational speed and/or infeasibility, number and choice of tuning parameters, restrictions on the number of “omic” features that can be selected, over-estimation of the number of relevant features and poor predictive performance are some of the noteworthy limitations when regularization is used on more general survival models that account for NPH or when a completely model-free method (such as SVM, random forests etc.) is used. Unlike the PH model, the AFT is built on the linear regression model for censored survival data and is a viable alternative to it since it directly models survival time and, thus, has a simpler and more intuitive interpretation. More importantly, it allows crossing hazard and survival curves, a useful property for modeling large-scale “omics” data with tens of thousands of features. In this paper, we adopt a more pragmatic approach for initially identifying the number of relevant features in a data set by simultaneously utilizing model significance and model fit based on different criteria. In addition, we adjust for potential confounders such as age of diagnosis and stage of disease with the goal of further eliminating spurious features. Such supervised marginal screening ensures that each feature selected for inclusion in the development of a prediction model actually fits the model of interest *and* has a statistically significant effect on survival.

Let *Y* denote the survival time of a typical subject in the study, the length of time entry into the study until a prescribed endpoint is attained. This endpoint may be the onset of a disease or event associated with it, or death itself. In addition, we let *C* be the duration of observation of the subject, i.e. the time from entry into the study until removal. The random variable *C* is referred to as a censoring variable. In general, we assume that both *Y* and *C* are non-negative random variables of which only the first one to occur is observed. Thus, an observation consists of the pair (*T, δ*), where *T* = min(*Y, C*) and *δ* = *I*(*T* = *Y*). We also have data on *p* covariates from each subject. It is assumed that censoring is non-informative, i.e. the survival time *Y* and the censoring mechanism *C* are independent, and that the covariates do not provide information about the censoring time *C*. Survival data usually consists of *N* samples, each containing the triple (*T*_*i*_, *δ*_*i*_, **z**_*i*_) for *i* = 1,…, *n*, where **z**_*i*_ = (*z*_*i*1_,…, *z*_*ip*_) is the covariate vector or profile of the *i*-th subject, *T*_*i*_ is the survival time if *δ*_*i*_ = 1 and it is the right censored time if *δ*_*i*_ = 0. The AFT model postulates a log-linear relationship between time and covariates given by,

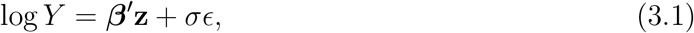

where *β′* is a vector of regression coefficients, **z** is the vector of covariates, *σ* is a scale parameter and *є* is the error term whose distribution is either pre-specified or is left completely unspecified, thus resulting in parametric or semi-parametric versions of the model (Kalbfleisch & Prentice, 2002; Jin et al., 2006). The intercept can be absorbed into *β′* and in the semi-parametric version, *σ ≡* 1 without loss of generality. Its log-linear form enables the measurements of the direct effect of features on survival time instead of the hazard; moreover, the regression coefficients can be interpreted in a similar fashion to that of multiple linear regression. In this model, effect size is measured as the ratio of expected survival times between two groups, say, patients exhibiting low and high expression of a particular feature or a set of features. In the clinical setting, it quantifies the effect of a feature on the expected duration of illness for a patient. This has lead many prominent statisticians, most notably Sir D. R. Cox, to observe that the AFT model and its estimated regression coefficients to have a rather ‘direct physical interpretation’ (Reid, 1994). Moreover, it is well-known that the PH and AFT models cannot simultaneously hold except in the case of extreme value error distributions. Therefore, the AFT model assumptions can hold when the PH model assumptions fail.

The semi-parametric AFT (*sAFT*) model is particularly attractive due to its distribution-free nature. Rank-based inference for this model is described in Jin et al. (2003), and regularized estimation is described in Cai et al. (2009) for high-dimensional data. An iterative solution has been developed to estimate the regression parameters (Jin et al., 2006). This procedure is based on the least-squares principle while accounting for censoring; however, it is computationally slow which can be problematic in the high-dimensional setting. The scope and applicability of AFT models can be significantly broadened by use of the generalized F distribution (*GenF*) (Ciampi et al. (1986) and more recently by Cox (2008)). *GenF* has the form seen in equation (4.4). Estimation for this model is based on maximum likelihood and, thus, offers a flexible and computationally efficient alternative to the *sAFT* model. Although *GenF* spans a variety of well known and lesser known models that are appropriate for modeling survival data, it has received little recognition in the literature. Its benefit lies in its umbrella structure and it embeds the generalized gamma (which includes Weibull, exponential, gamma and log-normal models), generalized log-logistic (which includes log-logistic models), F and Burr-type distributions. Other models such as the Maxwell-Boltzmann, generalized normal, half-normal, Chi and Raleigh are also members of this family among others. Thus, *GenF* provides a flexible approach to modeling patient survival in conjunction with large-scale “omics” data. There are several advantages to using *GenF*. As alluded to in *§*1, the AFT model intersects with the PH and PO models when the underlying data distribution is Weibull and log-logistic, respectively. The Weibull model with its monotonic hazard function and the log-normal model due to its mathematical intractability in dealing with censored observations offer limited potential for modeling survival data. Although the log-logistic model is similar in shape to the log-normal, its non-monotonic hazard function allows hazard curves to converge with time thereby incorporating a particular type of NPH and making it suitable for modeling cancer survival. It can be used if the course of the disease is such that mortality peaks after a finite time period and then slowly declines.

We motivate the utility of the AFT model for our problem using three data sets from large-scale cancer genomic studies that are detailed in *§*7.1. In this analysis, we fit semiparametric PH, PO and AFT models to each feature, after adjusting for clinical covariates such as age at diagnosis and stage of cancer, and evaluate their goodness-of-fit (GOF) using appropriate methods (Therneau & Grambsch, 1994; Martinussen & Scheike, 2006; and Novak, 2010). For each model, the *q*-value method was used to account for multiple testing (Storey & Tibshirani, 2003). The goal is to identify features that exhibit some form of NPH, thus demonstrating the need for alternatives to the PH model and, in particular, providing the rationale for a flexible model like AFT.

The results are summarized in Table 1 where A and B refer to sets of features for which the PH and PO model do not fit, respectively, and C refers to the set of features for which the AFT model fits, at the 5% significance level. Typically, we observe that there is a large number of features for which the PH or PO model does not fit across all data sets. More importantly, in each data set there is a significantly large fraction of features for which the AFT model fits (median of 97%). After correction for multiple testing, these observations are further corroborated by the corresponding *q*-values which indicate that the AFT model provides a good fit. The intersections of these sets is particularly revealing where we observe that the AFT model fits a large fraction of features for which the PH or PO model do not provide a good fit (median of 95%). Thus, it would be beneficial to develop methods based on the more general AFT model, which overlaps with the PH and PO models, due to its inherent ability to account for crossing hazards.

**Table 1:**
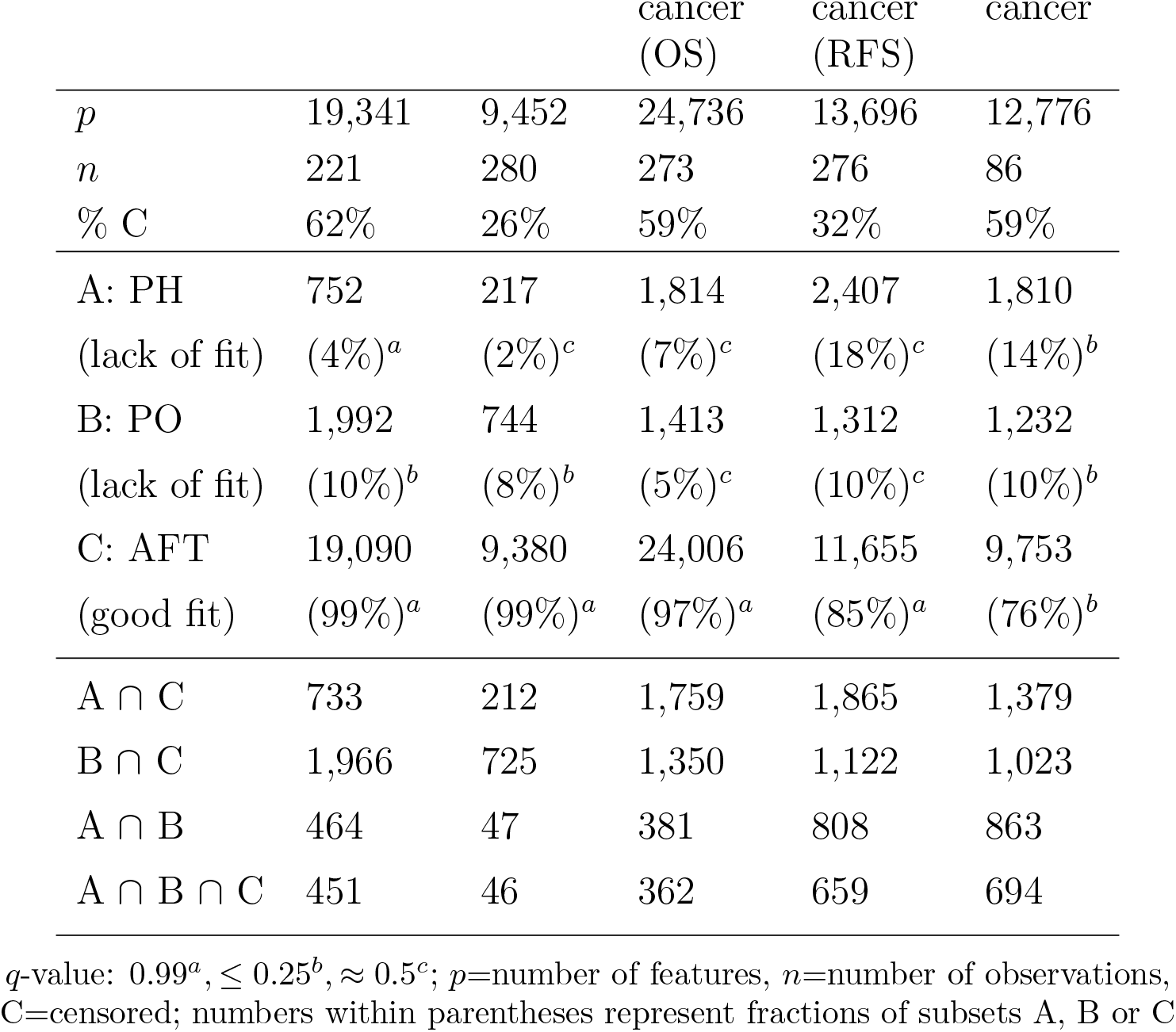
Summary of model fits

| Data Set | HNSCC | GBM | Ovarian<br>cancer<br>(OS) | Ovarian<br>cancer<br>(RFS) | Oral<br>cancer |
| --- | --- | --- | --- | --- | --- |
| $p$ | 19,341 | 9,452 | 24,736 | 13,696 | 12,776 |
| $n$ | 221 | 280 | 273 | 276 | 86 |
| % C | 62% | 26% | 59% | 32% | 59% |
| A: PH | 752 | 217 | 1,814 | 2,407 | 1,810 |
| (lack of fit) | (4%)<sup>a</sup> | (2%)<sup>c</sup> | (7%)<sup>c</sup> | (18%)<sup>c</sup> | (14%)<sup>b</sup> |
| B: PO | 1,992 | 744 | 1,413 | 1,312 | 1,232 |
| (lack of fit) | (10%)<sup>b</sup> | (8%)<sup>b</sup> | (5%)<sup>c</sup> | (10%)<sup>c</sup> | (10%)<sup>b</sup> |
| C: AFT | 19,090 | 9,380 | 24,006 | 11,655 | 9,753 |
| (good fit) | (99%)<sup>a</sup> | (99%)<sup>a</sup> | (97%)<sup>a</sup> | (85%)<sup>a</sup> | (76%)<sup>b</sup> |
| $A \cap C$ | 733 | 212 | 1,759 | 1,865 | 1,379 |
| $B \cap C$ | 1,966 | 725 | 1,350 | 1,122 | 1,023 |
| $A \cap B$ | 464 | 47 | 381 | 808 | 863 |
| $A \cap B \cap C$ | 451 | 46 | 362 | 659 | 694 |

$q$ -value: $0.99^a, \leq 0.25^b, \approx 0.5^c$ ; $p$ =number of features, $n$ =number of observations, C=censored; numbers within parentheses represent fractions of subsets A, B or C

## 4 Continuum power regression for large-scale “omics” data

We develop analytical methods for large-scale “omics” data using continuum power regression (CPR) - a unified framework for supervised dimension reduction - in conjunction with the AFT model. CPR embeds a spectrum of regression methods into a single framework that includes well known methods such as ordinary least squares (OLS), partial least squares (PLS) and principal components regression (PCR) as special cases. Stone & Brooks (1990) first proposed continuum regression (CR) and showed that OLS, PLS and PCR differed only in the target quantity being maximized in the process of extracting latent components that are linear combinations of these high-dimensional covariates. CR aims to maximize a quantity that includes the variation in covariates as well as the correlation of response with covariates, the relative proportions of which are controlled via a single parameter *γ*. At the extremes of this continuum, OLS maximizes correlation and PCR extracts orthogonal components by maximizing variance, while PLS lies in-between and maximizes the covariance between response and covariates. The numerical instability suffered by OLS due to multi-collinearity and high-dimensionality are circumvented by the unsupervised and supervised approaches provided by PCR and PLS, respectively, while the choice of *γ* provides further modeling flexibility.

Given an *n* × *p* matrix **Z** of predictors and an *n*-vector **t** of quantitative responses of an outcome, the objective function for constructing reduced components in CR can be expressed in terms of the objective functions for OLS (correlation, *R*^2^), PLS (covariance, *Cov*) and PCR (variance, *V ar*) as

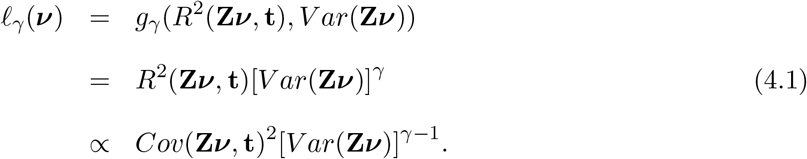

The optimization criterion is max_||***ν***||=1_ 𝓁_*γ*_ (***ν***) subject to ***ν***′***Sν***_*j*_ = 0, *j* = 1,…, *K*, where **S** = **Z**′**Z** is the covariance of **Z**, the columns of ***ν*** are weight vectors, *γ* ≥ 0 and *K* is the number of components. CR reduces to OLS (*γ* = 0), PLS (*γ* = 1) and PCR (*γ* → ∞) and can be shown to be closely related to ridge regression (Sundberg, 2002).

CPR is a variant of CR that is defined by the algorithm and not as the solution to the optimization problem in equation (4.1). In CPR, the PLS estimate ***τ** ∝* **ZZ**′**t** is generalized to ***τ** ∝* (**ZZ**′**t**)^*γ*^ for *γ* ≥ 0 where **Z** is modified into its powered version **Z**^(*γ*)^ via the SVD of **Z**, i.e., **Z**^(*γ*)^ = *UL*^*γ*/2^*V*′. CPR simplifies similar methods by requiring only one SVD after which standard PLS can be applied to **Z**^(*γ*)^, thus significantly improving computational speed and ease of interpretation (de Jong et al., 2001; Lorber et al., 1987). CPR coincides with CR for the special cases (*γ* = 0, 1 and ∞). It has been suggested that the continuity parameter *γ* and dimensionality *K* can play similar roles (Stone & Brooks, 1990; Frank & Friedman, 1993). There is also evidence to suggest that it is sufficient to consider only the three important special cases (OLS, PLS and PCR) and that the continuum may be unnecessary in CR under certain conditions (Chen & Cook, 2010). Given the fact that at three points CR is identical to CPR and that *K* and *γ* have similar effects, the simplicity, modeling flexibility and speed of CPR confer significant advantages over CR. In general, PLS requires fewer components than PCR; this is because the components from the latter need not necessarily be correlated with time-to-event, whereas all PLS components must be. PLS may be regarded as a compromise between OLS and PCR.

### 4.1 The CPR-AFT model

For a given application, CPR has the potential to offer insight into the underlying model. The ability of AFT to incorporate crossing hazard curves offers unparalleled flexibility for modeling large-scale “omics” data. As both CPR and AFT fall within the linear models framework, it seems natural to consider a hybrid model that combines their strengths. By combining AFT with CPR in a two-step procedure, we develop supervised dimension reduction methods jointly referred to as (A)CPR-AFT which include CPR-AFT and Adjusted CPR-AFT or ACPR-AFT, that adjusts for censored observations. (A)CPR-AFT represents a powerful array of solutions for this problem and enables identification of an “omic” profile that is predictive of a patient’s response to a specific treatment under a variety of scenarios encountered in practice.

(A)CPR-AFT has a distinct advantage over other methods in the literature because it directly addresses the three main issues with the application of survival analysis to “omics” data. First, it addresses the issue of high-dimensionality using CPR, reducing the number of “omic” features into a smaller number of CPR components that are linear combinations of these features. Second, it addresses the issue of NPH by using the AFT model, a model that does not assume PH but partly overlaps with the PH model. Lastly, it addresses the issue of censoring by imputing the censored observations using the extracted CPR components and the fitted AFT model. In the literature survey in *§*2 we noted other dimension reduction methods and, while some utilized either PLS or AFT, none of the methods addressed the issue of censoring directly. A large number of published large-scale genomic studies with censored survival outcomes seem to indicate that the proportion of censored observations is in the 60-80% range. Examples of such studies can be found in The Cancer Genome Atlas (TCGA) Network (http://cancergenome.nih.gov/), Gene Expression Omnibus (GEO) (https://www.ncbi.nlm.nih.gov/geo/) and Rouam et al. (2011), among others. Thus, in this application, having a method like ACPR-AFT that adjusts for censored data is not only beneficial but also desirable (Spirko, 2017). Furthermore, we explore the utility of CPR coefficients, *ω*, which are computed as *ω* = ***ν*****c**′ where the columns of ***ν*** are weights vectors and **c**′ are the loadings, in developing a survival prediction model and for feature ranking and selection.

#### 4.1.1 Supervised extraction of CPR components

The first step in the implementing the CPR-AFT model is to apply CPR which finds weight vectors, columns of ***ν***, such that the linear combinations **Z**^(*γ*)^***ν*** maximize the objective function 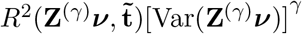 where 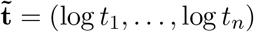 are the log transformed observed event times subject to the constraints outlined earlier. Here, **Z**^(*γ*)^ is found via the spectral decomposition of **Z**; after this step, standard PLS can be applied to **Z**^(*γ*)^. Let 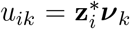 *i* = 1,…, *N*, *k* = 1,…, *K*, denote the linear combinations selected by CPR where 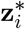 denotes the *i*^*th*^ row of **Z**^(*γ*)^ and ***ν***_*k*_ denotes the *k*^*th*^ column of ***ν***. These represent the CPR components, and the number of components *K* < *p* is chosen based on leave-one-out cross validation (LOOCV) to minimize the predicted residual sum of squares (PRESS) statistic. We use the reparametrization *γ ≡ α*/(1 − *α*) where *α* values of 0, 1/2 and 1 correspond to OLS, PLS and PCR, respectively. In subsequent sections, we outline how (A)CPR-AFT can be used to select *K*, the optimal number of components, and *α*, the CPR parameter. PLS is an important special case of CPR which maximizes the covariance between **Z*ν*** and log(*t*) and has been discussed by Devarajan et al. (2010) within this context.

#### 4.1.2 Fitting the AFT model to CPR components

The next step in the (A)CPR-AFT methods involve fitting the AFT model in equation (3.1). We propose flexible parametric and distribution-free versions of (A)CPR-AFT using the *GenF* and *sAFT* models, respectively. The flexible parametric AFT model, *GenF*, is given by

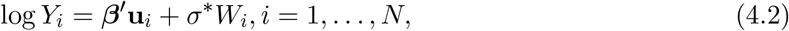

where *Y*_*i*_ is the survival time for the *i*-th subject, 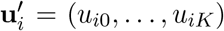 is a (*K* + 1) vector for the *i*-th subject, *β* = (*β*_0_, *β*_1_,…, *β*_*K*_) is the (*K* + 1) vector of unknown regression parameters, *W*_*i*_ are independent error terms with a common distribution *F*_*W*_ ∼ *GenF* and *σ*\* is the scale parameter. In this setting, *u*_*ik*_, *k* = 1,…, *K* represent the *K* CPR components for the *i*-th subject, *i* = 1,…, *N*. The semi-parametric AFT model, *sAFT*, has the form

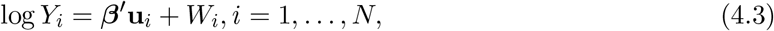

where the terms are as defined in equation (4.2). Here, *W*_*i*_ are independent error terms with unknown distribution *F*_*W*_. Given a feature expression vector **z** and PLS component **u**, using equation (4.2) the survival function of *Y* is given by

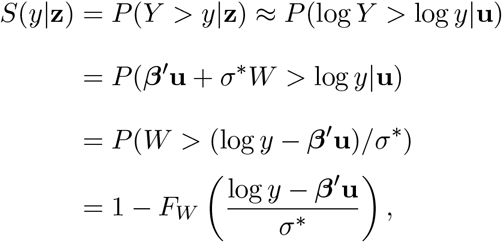

and is estimated by replacing the unknown parameters with their maximum likelihood estimates.

### 4.2 The parametric and semi-parametric (A)CPR-AFT algorithms

Below, we outline the CPR-AFT and ACPR-AFT algorithms. CPR-AFT ignores censoring and treats those observations as complete while ACPR-AFT imputes censored observations using mean residual life based on available data. For a pre-specified CPR parameter *α*, ACPR-AFT facilitates efficient extraction of the optimal number, *K*, of CPR components as determined by PRESS and LOOCV. The survival prediction algorithm proposed in *§*5, based on (A)CPR-AFT, simultaneously allows the optimal choice of *α* to be chosen in addition to the optimal *K* and utilizes the CPR coefficients, *ω*, from the final model.

#### Algorithm 1 CPR-AFT

1. Choose the parameter *α* and a pre-specified range of values for *K*.
2. For each *K* in the range pre-specified in Step 1, compute the PRESS statistic based on LOOCV and choose the number of CPR components, *K*, that minimizes the PRESS statistic. Note that, in this approach, *K* remains fixed and is chosen independently of *α*.
3. Perform CPR using *α* chosen in Step 1 and *K* identified in Step 2 to obtain weight vectors ***ν***_*k*_, *k* = 1,…, *K*. These weight vectors are used to compute the CPR coefficients *ω* as described in *§*4.1.1.
4. Build the final model using *ω* as detailed in *§*5.

#### Algorithm 2 ACPR-AFT

1. Repeat Steps 1-3 of CPR-AFT (Algorithm 1).
2. Use uncensored data to obtain CPR components where the number of components, *K*, is chosen as specified in Step 2 of the CPR-AFT algorithm. For *GenF*, use these components as covariates for the model in equation (4.2) and obtain estimates for ***β*** and *σ*\*; and for *sAFT*, use these components as covariates for the model in equation (4.3) and obtain the estimate for ***β***. It is important to note that the components obtained in this step are used only to estimate ***β*** and/or *σ*\*.
3. Let *v*_*i*_ = log *t*_*i*_. Impute censored observations by estimating the mean residual life using observed data *v*_*i*_ by 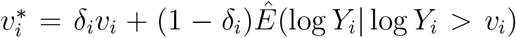, *i* = 1,…, *N*. Under the *GenF* model in equation (4.2), 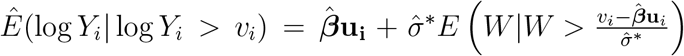, where 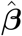 and 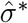 are obtained in Step 2 and *W* is the error term in equation (4.2). Under the *sAFT* model (4.3), 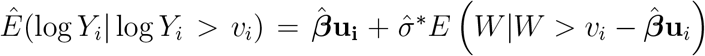, where 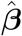 is obtained in Step 2 and *W* is the error term in equation (4.3). The calculation of this conditional expectation for *GenF* and *sAFT* models are outlined in *§*4.3 and *§*4.4, respectively. Here, ***u***_*i*_ are the CPR components obtained in Step 2 of CPR-AFT.
4. Use 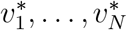 from Step 3 to construct new CPR components. The number of components, *K*, for the adjusted survival data is determined based on LOOCV such that the PRESS statistic is minimized. The weight vectors ***ν***_*k*_ corresponding to these new CPR components are used to compute the CPR coefficients *ω*.
5. Repeat Steps 1-4 for different choices of *α* and choose the optimal combination of (*α, K*) that minimizes the PRESS statistic.
6. Retain the CPR coefficients *ω* corresponding to the optimal choice of (*α, K*) in Step 5 to build the final model. This approach is outlined in detail in *§*5.

### 4.3 A flexible parametric approach to (A)CPR-AFT

In equation (4.2), if *Y* ∼ *GenF* then the density of can be written as

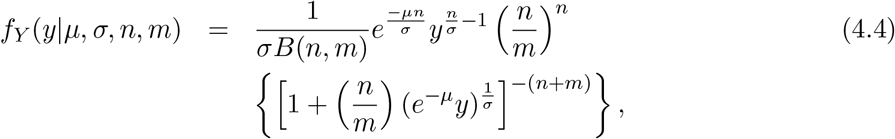

where *y* > 0. As shown in Ciampi et al. (1986) and Cox (2008), this model has an umbrella structure that includes many special cases where choosing specific parameter values will result in a particular model of interest. Important special cases include the generalized gamma, Weibull (exponential), gamma, log-normal, log-logistic, and Burr-type distributions and are listed in Table 2. The hazard behavior of *GenF* for finite values of the parameters is described in Cox (2008) and clearly highlights the flexibility provided by this model in handling different hazard shapes. Here, we propose a generalization of (A)CPR-AFT using the *GenF* model, which encompasses many important models and is, therefore, a flexible alternative for modeling censored survival data in conjunction with large-scale “omics” data. To this end, we first obtain the density of 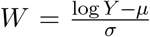 in equation (4.2) and use it to derive the expression for the conditional expectation 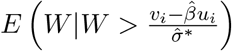 for this model in the ACPR-AFT algorithm. Using equation (4.4), the density of *W* can be derived to be

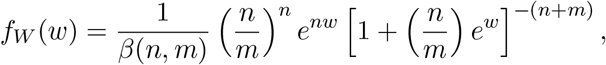

where −∞ < *w* < ∞. Using the density of *W*,

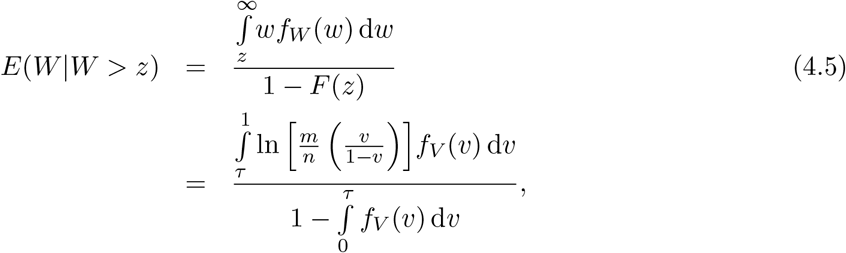

where 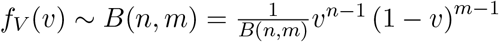, 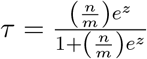 and 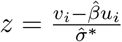

In CPR-AFT, *GenF* is used for model fitting in Step 3 while in ACPR-AFT, it is used to use to compute the expression in equation (4.5) in Step 3 and to fit appropriate models to estimate the parameters in Steps 2 and 5. The *GenF* model can be fitted using the R package flexsurvreg (R Core Team, 2018) and is described in Cox (2008). In addition to *GenF*, we are also interested in comparing its performance to important special cases such as the log-normal, log logistic, and Weibull. To obtain the conditional expectation, *E*(*W|W* > *z*), for these special cases, one just needs to replace the parameters in equation (4.5) with those listed in Table 2 for the respective model.

**Table 2:**
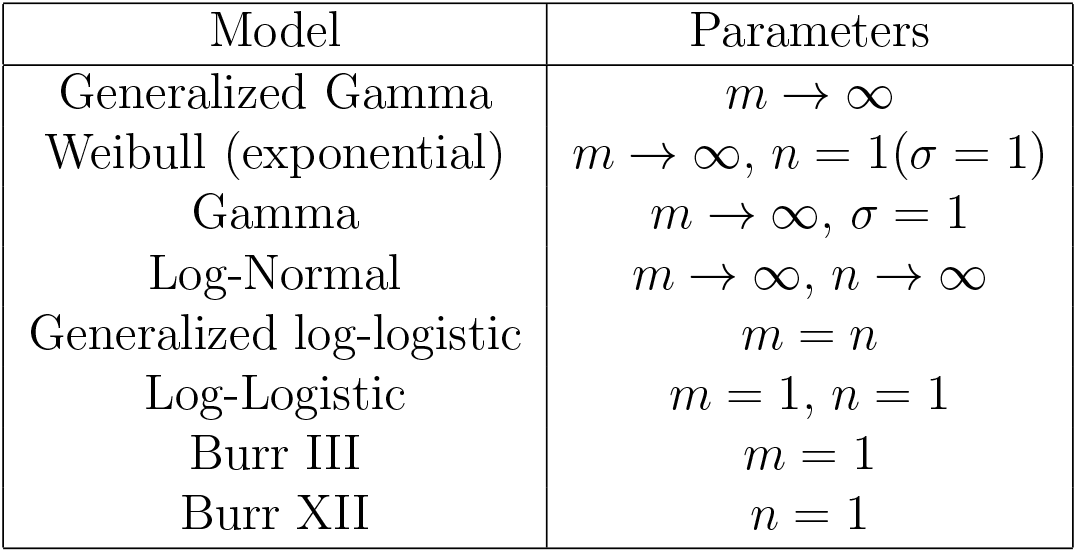
*GenF* Model: Some special Cases

| Model | Parameters |
| --- | --- |
| Generalized Gamma | $m \rightarrow \infty$ |
| Weibull (exponential) | $m \rightarrow \infty, n = 1(\sigma = 1)$ |
| Gamma | $m \rightarrow \infty, \sigma = 1$ |
| Log-Normal | $m \rightarrow \infty, n \rightarrow \infty$ |
| Generalized log-logistic | $m = n$ |
| Log-Logistic | $m = 1, n = 1$ |
| Burr III | $m = 1$ |
| Burr XII | $n = 1$ |

### 4.4 A semi-parametric approach to (A)CPR-AFT

Although the generalization based on *GenF* is parametric in nature, it offers tremendous modeling flexibility. Here, we further extend our (A)CPR-AFT approach using the *sAFT* model which has no distributional assumption for the error term. This is an attractive property as it does not force the choice of a specific model, thus providing more flexibility in the application of the method. The *sAFT* model has the form given in equation (4.2). Following Jin et al. (2006), the conditional expectation in Step 3 of ACPR-AFT is obtained as

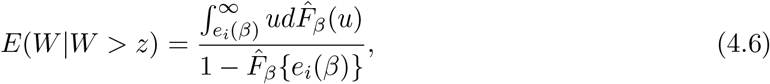

where 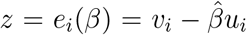 and 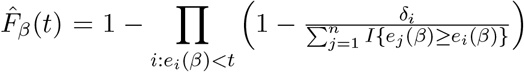 is the Kaplan Meier estimator of *F* based on {*e*_*i*_(*β*), *δ*_*i*_}. In CPR-AFT, *sAFT* is used for model fitting in Step 3 while in ACPR-AFT, it is used to compute the expression in equation (4.6) in Step 3 and to fit appropriate models to estimate the parameters in Steps 2 and 5. Fitting the semi-parametric AFT model is based on the Buckley-James (BJ) type estimator developed by Jin et al. (2006) and is implemented in the R package lss (R Core Team, 2018).

#### Merits of BJ Estimation

The BJ estimation method (Buckley & James, 1979) is an iterative least squares approach that is closely related to OLS without censoring and, thus, provides a more accessible interpretation to practitioners. It has been utilized in a variety of applications involving many areas such as medicine (Hammer et al., 2002), genetics (Bautista et al., 2008), astronomy (Steffen et al., 2006) and economics (Deaton and Irish, 1984; Calli and Wever bergh, 2009), and has been shown to be the preferred estimation approach in a comparison study (Wang & Wang, 2010). In contrast to methods that assume independence between the censoring mechanism and covariates, the BJ approach requires weaker assumptions and, in conjunction with boosting, has been shown to be superior to LASSO-type methods and to generate sparser models (Wang & Wang, 2010). It utilizes Kaplan-Meier estimates and is readily available in statistical software such as R (R Core Team, 2018); moreover, BJ estimation for the AFT model can be conveniently extended to describe more complex data structures with existing software, such as MART and MARS (Friedman, 1991; 2001). Hence, we use BJ estimation in the proposed algorithms.

## 5 Supervised dimension reduction

A natural approach to build a final model based on (A)CPR-AFT is using the *K* CPR components from ACPR-AFT as covariates in equation (4.2) and following Step 3 of CPR-AFT. It would result in an AFT model based on reduced components from ACPR-AFT. A prognostic index can be defined as *η* = **u***β*, where **u** is the *n* × *K* matrix whose columns contain the *K* CPR components and *β* is the *K*-vector of coefficients from this final AFT model fit. A major disadvantage of this approach is that component information is not available for new subjects and, therefore, it is not possible to develop a prediction model that can be used on future subjects with feature expression profiles. Recall that *ω*_*p*×1_ = ***ν***_*p*×*K*_ **c**′_*K*×1_ where the *K* columns of ***ν*** contain weight vectors and **c**′ contains the loadings associated with the *K* CPR components which are contained in the columns of **u**_*n*×*K*_. Hence, we propose an approach based on the CPR coefficients, *ω*, rather than directly using the reduced CPR components, **u**, for predicting the survival probability of a future subject whose feature expression profile is readily available. As shown in the next section, some important differences exist between this approach and the final AFT model discussed above with vastly different implications for prediction.

### 5.1 Developing a prognostic index

We use the CPR coefficients to devise an approach based on the weighted average of feature expressions and illustrate its utility in developing a survival prediction model. Using the vector of CPR coefficients, *ω*, for the *p* features from a particular model of interest and the *n* × *p* feature expression matrix, **Z**, the weighted average, *η*, is calculated as

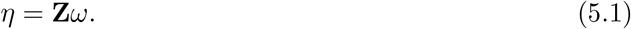

This results in an *n*-dimensional vector, which we call the *prognostic index (PI)*, where each element represents a subject’s weighted average feature expression. In the calculation of *η*, a heavier weight is placed on features deemed significant and in our approach, we calculate *η* using the subset of features determined by marginal screening procedures (see *§*5.2 for details). It is worth noting that *PI* represents the predicted (log) survival times and, thus, enables the development of a prediction model using (A)CPR-AFT as outlined in the next section.

### 5.2 Predictive modeling using (A)CPR-AFT

We develop a survival prediction algorithm using the CPR coefficients *ω* from (A)CPR-AFT, separately for *GenF* and *sAFT*, by adopting a flexible approach that simultaneously chooses the optimal *α* in addition to the optimal *K*, the number of CPR components. The proposed approach utilizes (A)CPR-AFT for several choices of *α* that represent a variety of scenarios: the midpoint of the trajectory from OLS and PLS (*α* = 0.25), PLS (0.5), the midpoint of the trajectory from PLS to PCR (0.75) and PCR (0.95). Since *p* ≫ *n*, OLS (*α* = 0) is not a useful option in our application and PCR (*α* = 1) requires a value of *α* close to 1 in order to avoid numerical instability. In this approach, the final model is chosen based on the optimal (*α, K*) combination that results in the smallest PRESS using LOOCV after applying (A)CPR-AFT, and the corresponding CPR coefficients *ω* are used to develop the *PI* for evaluating the performance of this prediction model.

In CPR-AFT, there is no way of choosing an optimal *α* because it plays no role in the selection of *K* and is only used after *K* is chosen to run CPR. Therefore, an optimal (*α, K*) combination cannot be chosen because Step 2 (choosing *K* using PRESS) does not depend on *α*. *K* would remain the same even if Steps 3 and 4 of CPR-AFT are repeated for different choices of *α* where each *α* would yield a different survival prediction model, and models corresponding to various choices of *α* would have to be evaluated and compared separately. On the other hand, an optimal combination can be chosen in ACPR-AFT because each pre-specified *α* directly impacts the adjustments. By comparing the PRESS statistics after adjustments are made (step 4 of ACPR-AFT) for different choices of *α*, the optimal (*α, K*) combination can be selected. Thus, ACPR-AFT has a significant advantage over CPR-AFT because it adjusts for censored observations. For these reasons, we consider two different unadjusted methods in our comparisons, one each based on the chosen value of *α* from *GenF* and *sAFT* in ACPR-AFT.

Prior to the application of (A)CPR-AFT, supervised marginal screening procedures were used to narrow down the number of features. These methods ensure that features used for prediction demonstrate an association with survival, at the univariate level, after adjusting for potential confounders such as age of diagnosis and stage of disease. An added benefit of such pre-filtering is that it significantly reduces computation time. Supervised marginal screening was performed to select (i) features that fit the *sAFT* model and had a statistically significant effect on survival (sAFT) or (ii) features that had a significant effect on survival using concordance regression (CON) (Dunkler et al., 2010), at the 0.05 significance level. Once a subset is selected, (A)CPR-AFT is applied and the optimal (*α, K*) is chosen. The CPR coefficients, *ω*, are retained for the adjusted (ACPR-AFT) and unadjusted (CPR-AFT) methods (based on *GenF* and *sAFT*) and used to predict the logarithm of survival time for each subject given their feature expression profile **Z** using the prognostic index, *PI* = *η* = **Z***ω*. We use italicized notation (*sAFT* or *GenF*) to denote the particular method associated with ACPR-AFT while sAFT is used to denote the marginal screening method.

The following cross validation approach is used to build and evaluate the prediction models. The data is first split into training and test sets roughly in a 2:1 ratio, where *ω*_*tr*_ represents the vector of CPR coefficients corresponding to the training set and is used to predict the logarithm of survival time in the test set. Thus, *PI* = **Z**_*te*_*ω*_*tr*_, where **Z**_*te*_ is from the test set, and the model is evaluated for prediction accuracy. We utilize the following measures of prediction accuracy to evaluate and compare the predictive performance of ACPR-AFT, using *GenF* or *sAFT* models, to the unadjusted CPR-AFT approach: (i) *R*^2^, the fraction of variation that is explained by the *K* CPR components in the final (A)CPR-AFT model, (ii) Mean Squared Error, 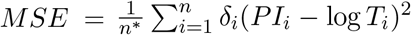 where *PI*_*i*_ is the prognostic index for the *i*^*t*^*h* subject, 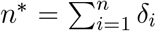 and *δ*_*i*_ = 1 implies the event was observed; *MSE* is calculated for both the training set, *MSE*_*TR*_, and the test set, *MSE*_*TE*_, and (iii) area under the time-dependent receiver operating characteristic curve (*AUC*) which quantifies a method’s ability to predict survival at varying time points such as 2, 3 or 5 years and is implemented in the R package survivalROC (Haegerty et al., 2000; R Core Team, 2018). An *AUC* close to 1 indicates better prediction accuracy. In summary, the survival prediction algorithm involves the following steps:

#### Algorithm 3 Survival Prediction Algorithm

1. Use supervised marginal screening to filter features using sAFT or CON, as outlined above.
2. Randomly split the filtered data set into training (67% of subjects) and test sets (33% of subjects).
3. Apply (A)CPR-AFT (*GenF* and *sAFT*) to the training set using *α* = (.25, .5, .75, .95).

- Choose optimal (*α, K*) combination.
- Retain the CPR regression coefficients, *ω*_*tr*_.
4. Use *ω*_*tr*_ from Step 3 to predict (log) survival times in the test set, i.e., calculate *PI* = **Z**_*te*_*ω*_*tr*_, where **Z**_*te*_ is from the test set.
5. Evaluate the prediction models using the measures of prediction accuracy outlined above.
6. Repeat Steps 2-5 25 times. Median values of the prediction accuracy measures from step 5 are reported.

## 6 Application to simulated data

### 6.1 Simulation schemes

We considered two different simulation schemes to generate artificial survival and feature expression data sets based on the approach outlined in Dunkler et al. (2010). In order to account for various types of hazards, survival times *Y*_*i*_, *i* = 1,…, *n*, were generated from each of 5 different models specified as follows: standard log-normal LN (*µ* = 0, *σ* = 1); log-logistic LL1 (*α*_1_ = 2, *λ*_1_ = 2, *λ*_2_ = 4) and LL2 (*α*_1_ = 3, *α*_2_ = 4, *λ*_1_ = 1, *λ*_2_ = 2); and Weibull W1 (*α*_1_ = 1, 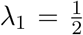) and W2 (*α*_1_ = 3, *α*_2_ = 2, *λ*_1_ = 1, 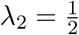), where LL1 and W1 refer to the respective models where the shape parameters are the same but the scale parameters differ, and LL2 and W2 refer to the respective models where both the shape and scale parameters differ. We use a more informed approach that is broader in scope compared to that of Dunkler et al. (2010), who only considered W1 in their simulations. Here, LN, LL2 and W2 cases are of particular interest because of their ability to simulate crossing hazards. To simulate censoring, we drew random samples with uniform follow-up times *C* from *U* (0, *τ*) and defined the observed survival time as *T* = min(*Y, C*) with censoring indicator *δ* = *I*(*T* = *Y*). We chose *τ* to get censoring proportions of 33, 67% and 80%.

For each model, we simulated censored survival times and feature expression data for *N* = 200 subjects and *p* = 5000 mock features where feature expression is linked to survival time based on the logarithm of the hazard ratio (HR), *β*_*g*_(*t*) = *β*_0_ log(*HR*). Feature expression data was generated from the standard normal model. Following Klein and Moeschberger (2003), log(*HR*) was calculated based on the respective model of interest. For LN, we used *β*_*g*_(*t*) = *β*_0_(*t*^2^ − 1) to simulate crossing hazards similar to what was done in Dunkler et al. (2010). Then, *β*_0_ was chosen so that only the first 400 features were assumed to have an effect on survival time, with 200 having a large effect and 200 having a small effect. In Scheme 1, we adopt a univariate approach where feature expression is linked to survival one feature at a time, and in Scheme 2 we adopt a multivariate approach that incorporates correlations between features. More details on these steps can be found in Dunkler et al. (2010).

### 6.2 Evaluation of methods

In SI *§*1.1, we discuss feature ranking and selection methods using components extracted from (A)CPR-AFT. We evaluated their performance for (i) *GenF* versus its special cases and (ii) *GenF* and *sAFT* -based ACPR-AFT versus unadjusted CPR-AFT under different data generating mechanisms and censoring fractions for each simulation scheme. These results are summarized in SI Tables 1-6 and Figures 1 & 2. In addition, the survival prediction algorithm proposed in *§*5.2 was evaluated using *GenF* or *sAFT* -based ACPR-AFT and compared with unadjusted CPR-AFT for each simulation scheme. Details are provided in the SI *§*1.2 and results are summarized in SI Tables 7-12. Overall, our simulation studies establish the superiority of *GenF* or *sAFT* -based ACPR-AFT under a variety of data generating mechanisms encountered in practice.

## 7 Application to high-throughput “omics” data

We demonstrate the utility of (A)CPR-AFT in supervised dimension reduction and developing a survival prediction model using the following publicly available data sets in cancer genomics. These data sets are described in detail in SI *§*2.

### 7.1 Data sets

- Head & Neck squamous cell carcinoma (HNSCC): Published by TCGA and contains survival data and RNA sequencing gene expression profiles for 221 subjects with HNSCC.
- Glioblastoma (GBM): Published by TCGA and contains survival data and methylation profiles for 280 tumor samples obtained using the Infinium HumanMethylation27 platform.
- Ovarian cancer: Published by Tothill et al. (2008) and contains Affymetrix gene expression profiles for 282 subjects and corresponding overall survival (OS) and recurrence-free survival (RFS) data.
- Oral cancer: Published by Saintigny et al. (2011) and contains survival data and gene expression profiles for 86 subjects obtained using the Human Gene 1.ST platform.

### 7.2 Extracting genomic components for predictive modeling

We illustrate the utility of (A)CPR-AFT for predictive modeling by applying the survival prediction algorithm to the HNSCC data set. Table 3 summarizes the results for each marginal screening method. In each case, the optimal value of *α* chosen by ACPR-AFT was used for the corresponding unadjusted method. A significant improvement in predictive performance is noted between each adjusted method (*sAFT* or *GenF*) and CPR-AFT for both screening methods in terms of at least two out of the three evaluation measures used, *R*^2^, *MSE* and *AUC*. Next, we illustrate the application of (A)CPR-AFT in supervised dimension reduction using the four genomic data sets outlined above for the special case *α* = 0.5 (PLS) where ACPR-AFT is based on *GenF* or *sAFT*. In each case, the optimal number of CPR components, *K*, was determined using the PRESS statistic based on LOOCV by considering ranks *k* = 2,…, 15. The results, summarized in Table 4, indicate that for all data sets *GenF* - and *sAFT* -based APCRAFT generally outperform the unadjusted method by explaining a higher proportion of the variation in the data at the chosen optimal *K* or by choosing a smaller optimal *K*. In some cases, ACPR-AFT is observed to choose at least as many components as the unadjusted method; however, even at the optimal *K* chosen by the unadjusted method (ovarian OS and oral) or at a lower rank (HNSCC, GBM, ovarian RFS and oral), ACPR-AFT explains a higher fraction of the variation compared to the unadjusted method (as shown in the last three rows of Table 4). For the GBM and ovarian RFS data sets, ACPR-AFT performs as well as the unadjusted method; this is likely due to the relatively small fraction (26% and 32%, respectively) of censored observations in these sets compared to others where censoring ranges from 59-62%. These examples suggest that it is possible to choose a more parsimonious model than that provided by ACPR-AFT while still explaining most of the variation in the data. Moreover, they highlight the utility of ACPR-AFT for handling censored data. The CPR coefficients, *ω*, and VIP, *ξ*, obtained from ACPR-AFT can also be used for feature ranking purposes as demonstrated by our simulations (see SI *§*1.1 for details).

**Table 3:** Prediction Summary (HNSCC Data)

| Filter<br>(# of features) | Measure | ACPR-AFT<br>*GenF* | ACPR-AFT<br>*sAFT* | CPR-AFT<sup>a</sup> |
| --- | --- | --- | --- | --- |
| sAFT<br>(2,336) | $R^2$ | 100 | 99.8 | 98.3 |
| | $MSE$ | 0.004 | 0.07 | 0.53 |
|  | AUC, t = 2 | 0.81 | 0.96 | 0.73 |
|  | AUC, t = 3 | 0.85 | 0.95 | 0.71 |
|  | AUC, t = 5 | 0.67 | 0.89 | 0.67 |
| CON<br>(2,254) | $R^2$ | 99.0 | 99.1 | 91.3 |
| | $MSE$ | 0.95 | 0.62 | 1.18 |
|  | AUC, t = 2 | 0.69 | 0.92 | 0.69 |
|  | AUC, t = 3 | 0.70 | 0.93 | 0.69 |
|  | AUC, t = 5 | 0.67 | 0.9 | 0.66 |

*a*: based on optimal $\alpha$ chosen by the corresponding ACPR-AFT method used. *GenF* and *sAFT* chose the same optimal $\alpha$ .

**Table 4:** (A)CPR-AFT - Summary of Results

| Method | HNSCC |  | GBM |  | Ovarian OS (RFS) |  | Oral |  |
| --- | --- | --- | --- | --- | --- | --- | --- | --- |
| | $K$ | $R^2$ | $K$ | $R^2$ | $K$ | $R^2$ | $K$ | $R^2$ |
| *GenF*<sup>†</sup> | 8 | 100% | 5 | 100% | 2 (13) | 96% (100%) | 9 | 100% |
| *sAFT*<sup>†</sup> | 8 | 100% | 5 | 100% | 2 (15) | 97% (100%) | 5 | 97% |
| CPR-AFT | 12 | 100% | 5 | 100% | 2 (15) | 95% (100%) | 5 | 84% |
|  | 2 | 92% | 2 | 93% | * (2) | * (94%) | 2 | 79% |
| *GenF*<sup>†</sup> | 2 | 96% | 2 | 93% | * (2) | * (94%) | 5(2) | 97% (86%) |
| *sAFT*<sup>†</sup> | 2 | 98% | 2 | 94% | * (2) | * (95%) | 5 (2) | 97% (84%) |

<sup>†</sup> ACPR-AFT; \* $K = 2$ chosen by CPR-AFT and ACPR-AFT

### 7.3 Interpreting the prognostic index

The prognostic index, *P I* = *η* = **Z***ω* can be interpreted as a linear predictor and is particularly relevant within the context of (A)CPR-AFT which combines two linear models. The predicted survival curves obtained from *GenF* - and *sAFT* -based ACPR-AFT using *η*, plotted in SI Figure 3 for each data set, thus illustrate the use of CPR coefficients, *ω*, in building a prediction model. In each case, the corresponding model under consideration (*GenF* or *sAFT*) was statistically significant at the 5% significance level. Furthermore, GOF tests performed using the methods outlined in *§*3 revealed a good fit for the PH, PO and AFT models across different methods and data sets. The only exceptions to this were a lack of fit for the PH model for both *sAFT* - and *GenF* -based ACPR-AFT for the ovarian OS data set which indicates an overall time-varying effect due to features selected by the model. While a weighted average using VIP, *ξ*, could serve as a prognostic index, it does not directly predict survival time and, hence, its interpretation is unclear (see SI *§*1 for details). These examples thus serve to illustrate the utility of *η* in elucidating the relationship between feature expression and patient survival and to account for time-varying effects of features.

### 7.4 Evaluating the prediction algorithm

The survival prediction algorithm outlined in *§*5.2 was evaluated using the four data sets. After marginal screening, the prediction algorithm was applied separately to each training and test set. Median values of different measures of prediction accuracy, based on 25 random training and test sets, are summarized in SI Tables 13-17. Overall, less variation in these measures was observed for ACPR-AFT across the training and test sets and for both filters used compared to CPR-AFT; in particular, *GenF* showed much less variation in *R*^2^ and MSE compared to *sAFT* while the opposite effect was observed for AUC (data not shown). In each run, the optimal (*K, α*) combination was chosen as outlined earlier for ACPR-AFT. Since CPR-AFT does not involve choosing *α* based on adjustment (as in Step 4 of ACPR-AFT), the optimal *α* chosen by *GenF* - and *sAFT* -based ACPR-AFT is used for comparison purposes.

The fraction of variation in the data explained by the CPR components extracted by a particular model, quantified by *R*^2^, are significantly higher for *GenF* - and *sAFT* -based ACPR-AFT compared to the unadjusted CPR-AFT. Since computation of *R*^2^ requires model fitting, it is relevant only to the training set. Not surprisingly, *MSE* is generally larger for the test set compared to the training set across both marginal screening methods and data sets. However, for both the training and test sets, we consistently observe smaller *MSE* for ACPR-AFT methods compared to CPR-AFT across all four data sets which indicates that both *GenF* and *sAFT* result in more accurate predictions. In particular, a significant reduction in *MSE* between ACPR-AFT and CPR-AFT is observed for the oral test sets, ranging from 58-91% across both filters. For the ovarian RFS and HNSCC test sets, CON (18%) and sAFT (26%) filters result in maximum reduction, respectively.

In addition, we examine the AUCs calculated for 2, 3 and 5 year survival. Once again, we note that the training set AUCs are larger than those of the test set in each case. More importantly, in both the training and test sets and across both filtering mechanisms, we observe larger AUCs for *GenF* and *sAFT* compared to the unadjusted methods. The improvements observed in the test sets are particularly relevant and are noteworthy for all data sets. For example, using the sAFT and CON filters, at *t* = 5 we observe an AUC range of 0.74-0.82 for *GenF* and *sAFT* for the ovarian OS test sets while the unadjusted methods range only from 0.68-0.73. The performance of the two filters was similar for the ovarian (OS & RFS) and HNSCC data sets and resulted in improvements up to 10% and 13% in AUC, respectively; for the GBM data set, however, CON provided substantial improvement in AUC of up to 15%. In addition, a statistically significant difference was generally observed between the AUCs from time-dependent ROC curves for ACPR-AFT and CPR-AFT.

## 8 Conclusions and discussion

In this paper, we proposed supervised dimension reduction methods for analyzing large-scale “omics” data in conjunction with censored survival outcomes. Our methods combine CPR - a unified framework that includes OLS, PCR and PLS as special cases - for dimension reduction with the AFT model - a censored linear regression model - for handling survival data, and offer distinct advantages relative to currently available approaches. The versatility afforded by the parametric (*GenF*) and semi-parametric (*sAFT*) versions of the AFT model and its partial overlap with the widely used PH and PO models allow a variety of time-varying feature effects to be incorporated. Moreover, both CPR and AFT fall within the linear models framework and the proposed hybrid model, (A)CPR-AFT, combines their strengths in a unique fashion that does not match any other available method. The umbrella structure of *GenF* provides a fully parametric, yet tremendously flexible, approach for modeling survival data while *sAFT* utilizes BJ estimation which has been shown to be a robust method. A particularly attractive characteristic of ACPR-AFT is its ability to account for censored observations common to studies with survival endpoints. Many large-scale “omics” studies involving survival outcomes of interest tend to contain a significant fraction of censored observations and an appropriate method for handling these incomplete observations has been lacking. The simulation results demonstrated the superior predictive performance of ACPR-AFT over CPR-AFT under a plethora of data generating mechanisms particularly as the fraction of censored observations increased, thus making it a practically useful tool for data analysis. These results were corroborated using publicly available data sets in cancer genomics where the performance of the proposed survival prediction algorithm was shown to improve significantly when censoring was accounted for in this manner.

The ability of the proposed methods to handle NPH within this context is unparalleled and it offers a robust and flexible approach for predictive modeling of a wide variety of large-scale “omics” data. The CPR coefficients and VIP play complementary roles and serve different purposes. The former is useful for computing the *PI* which was used to develop and evaluate the prediction algorithm while the latter was shown to be a superior measure for feature ranking. However, choosing an appropriate threshold for these measures is an important consideration and could form part of future work on this topic. When combined with an appropriate marginal screening method, this approach could serve as a useful feature selection tool by significantly reducing the number of relevant features in the prediction algorithm which could, in turn, not only improve its performance further but also help develop a feature signature that is predictive of survival. Regularization is another avenue for future research on ACPR-AFT that would aid in feature selection.

Furthermore, the proposed methods are broadly applicable to a variety of high-throughput “omics” data such as feature expression data arising from next-generation sequencing, allele-specific expression, methylation, microarrays and SNP arrays as well as large-scale data from proteomics, metabolomics and DNA copy number studies, many of which have been utilized in this study. There has been a recent surge in integrative “omic” analyses that simultaneously involve different data types as well as other quantitative outcome variables using publicly available data from repositories such as TCGA and GEO (Ramakodi et al., 2016; Li et al., 2013; Lawrenson et al., 2015). Within this context, the proposed unifying framework offers a robust platform for analysis and interpretation.

## Acknowledgement

The work of KD was funded in part through the NIH/NCI Cancer Center Support Grant P30 CA006927.

## Supplementary Information

### 1 Simulation Results

#### 1.1 Feature ranking and selection using (A)CPR-AFT

In this section, we discuss the utility of (A)CPR-AFT for feature ranking and selection by comparing the performance of (A)CPR-AFT algorithms based on log-normal (LN), log-logistic (LL), Weibull (W), *GenF* and *sAFT* models. Wold et al. (2002), Devarajan et al. (2010) and Mehmood et al. (2012) discuss the use of PLS coefficients, *ω*, and variable importance projection (VIP), denoted by *ξ*, as measures for feature ranking and selection in PLS. Note that the PLS coefficients arise as a special case of CPR when *α* = 0.5. For example, features could be ranked based on the absolute value of CPR coefficients, *ω*, which can take on values on the entire real line, or directly using the VIP, *ξ*, which is a non-negative quantity.

VIP accumulates the importance of each feature as reflected by the weight ***ν**_k_* from each component. Essentially, it is a measure of the contribution of each feature according to the variance explained by each component. The VIP value, *ξ*, for feature *j* is calculated as

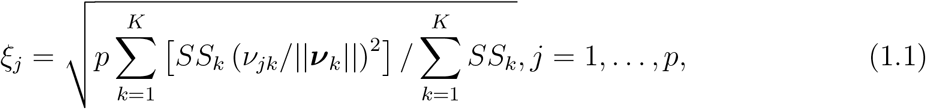

where *p* is the number of features, *K* is the number of components, ***ν***_*k*_ is the weight vector for the *k*-th component, and *SS*_*k*_ is the sum of squares explained by the *k*-th component. In other words, CPR produces *K* vectors of weights, each of which has *p* elements corresponding to the *p* features. In the VIP calculation, (*ν*_*jk*_/||***ν***_*k*_||)^2^ represents the importance of the *k*-th component. Thus, *ξ*_*j*_ is calculated for feature *j*, *j* = 1 … *p*, and then features are selected based on a threshold pre-specified by the user. A popular threshold is 1 (i.e., *ξ* > 1) and is discussed in Mehmood et al. (2012). Since the number of relevant features is pre-determined in simulations, a threshold is not relevant in our application where features are ranked separately based on decreasing VIP, *ξ*, and absolute value of *ω*. Although choice of a threshold is unclear in the analysis of real data, these measures can be used for feature ranking and selection under certain scenarios. For example, *ξ* could be used to rank features identified in the survival prediction algorithm proposed in §5.2, similar to how the corresponding *ω* is used to calculate *PI*. We plan to investigate choice of the *ξ* threshold in future work. In this paper, we utilize *ξ* only as a feature ranking tool as illustrated in our simulation studies. Our analyses using a variety of large-scale “omics” data sets showed that *α* = 0.5 resulted in overall better performance in terms of model fit and prediction accuracy; hence, we focus on this special case for the simulation studies. As mentioned earlier, high censoring appears to be a common theme in many large-scale “omics” studies involving censored survival outcomes (Bhattacharjee et al., 2001; Beer et al., 2002; Tothill et al., 2008; Saintigny et al., 2011; Rouam et al., 2011; TCGA Network, GEO); hence, this scenario is also of particular interest in the simulations. As evidenced in the following sections, our studies indicate that VIP is a more useful quantity for feature ranking but less useful for the purposes of developing a prediction model where the CPR coefficients play a significant role.

For each simulation scheme and censoring combination, 200 randomly generated data sets were created and assessed. The (A)CPR-AFT algorithm was applied to each simulated data set and mock features were ranked separately using the absolute value of PLS coefficients, *ω*, and VIP, *ξ*. In the remainder of this section, we will only use *ω* or *ξ* to denote these methods. For CPR-AFT, *ω* and *ξ* are calculated in Step 2 of the algorithm, and therefore, are not model specific. On the other hand, in ACPR-AFT these quantities are computed after adjusting censored observations in Step 4, thus taking the pre-specified model into account in the parametric version or in a completely distribution-free manner in the case of *sAFT*. In the parametric version, we considered *GenF* and three of its well-known special cases - LN, LL and W. For each method, the ranked lists were used to compute mean values of sensitivity, specificity, Youden Index (=sensitivity+specificity-1) (Youden, 1950) and area under the receiver operating characteristic (ROC) curve (AUC) across the 200 data sets. The purpose of this analysis is to compare the performance of (i) the two ranking methods, *ω* and *ξ*, (ii) *GenF* vs. its special cases in ACPR-AFT, and (iii) CPR-AFT vs. ACPR-AFT for *GenF* and *sAFT* models under both simulation schemes.

##### 1.1.1 Comparison of *GenF* and its special cases in ACPR-AFT

We examined the performance of *GenF* and its special cases in ACPR-AFT using AUC and the Youden Index for 33% and 80% censoring. SI Figure 1 depicts the ROC curves for 33% and 80% censoring for *ω* and *ξ* using simulation scheme 1. In all four cases - shown in panels (a)-(d) - we observe that *GenF* outperforms LN, W, and LL. The corresponding AUCs and Youden Indices are reported in SI Table 5. The AUCs for *GenF* are higher than its special cases in each situation, and the differences increase as censoring increases. We note that the unadjusted CPR-AFT results in the lowest Youden index and AUC in every scenario and performs significantly worse than ACPR-AFT using *GenF*. In addition, we observe that *ξ* is superior to the use of *ω* in terms of AUC and the Youden Index, thus indicating that VIP is a better tool for feature ranking and selection.

Next, we evaluated the performance using simulation scheme 2 for 33% and 80% censoring. Panels (a) and (b) of SI Figure 2 show the ROC curves for these censoring proportions, respectively, comparing the performance of *ω* and *ξ* for *GenF*. Similar to scheme 1, we observe that *ξ* significantly outperforms *ω*. Panels (c) and (d) of SI Figure 2 show the ROC curves for 33% and 80%, respectively, comparing the performance of *GenF* and CPR-AFT for *ξ*. Again, it is evident that *GenF* outperforms the unadjusted method. Although not shown, scheme 3 showed similar results for *GenF* compared to its special cases LN, LL and W. At each censoring level, *GenF* had the largest AUC and Youden index, similar to what was observed for scheme 1 in SI Figure 1 and SI Table 5. Since *GenF* outperformed its special cases, we will focus only on *GenF*, *sAFT* and the unadjusted method for the remainder of the paper. A very similar performance was observed across multiple simulated data sets and thus, results are reported only for a single, representative data set.

##### 1.1.2 Simulation Scheme 1

In this section, we evaluate the performance of *GenF* - and *sAFT* - based ACPR-AFT against unadjusted CPR-AFT using data simulated from scheme 1 as described in §5.1. SI Table 6 shows the AUCs using *ξ* as the feature selection tool. Across each scheme and censoring level, we observe that the proposed ACPR-AFT method (both *GenF* - and *sAFT* -based) have a clear advantage over CPR-AFT, and as the censoring increases, the differences in performance become stronger. In fact, CPR-AFT has the lowest AUC in every single scheme for the 67% and 80% censoring cases. Thus, we observe a clear benefit when imputing censored observations using ACPR-AFT. Next, we examine differences between *GenF* - and *sAFT* -based ACPR-AFT. Both *GenF* and *sAFT* perform similarly in many instances, but there are particular schemes where one outperforms the other. For example, AUCs of *GenF* are higher for the LN, LL1 and LL2 cases, and AUCs of *sAFT* are higher in the W1 and W2 cases. In SI Table 7, AUCs using *ω* are shown and observed to be lower than those of *ξ* in each case. Thus, *ξ* appears to be the optimal approach for feature selection; however, even when *ω* is used, *GenF* and *sAFT* outperform CPR-AFT.

Next, we focus our attention on the Youden index displayed in SI Table 8 for each method and censoring level using both *ω* and *ξ* as ranking methods. First, we note that Youden indices based on *ω* are lower than those based on *ξ* in every single case. Thus, as expected, *ξ* significantly outperforms *ω* in ranking features in every scenario. Furthermore, we observe that *GenF* - and *sAFT* -based ACPR-AFT outperform CPR-AFT in almost every scenario, and just as we had observed with the AUCs, the differences in performance become larger as censoring increases. Thus, we notice a clear improvement due to ACPR-AFT. *GenF* and *sAFT* perform similarly in many cases, but as indicated by the AUC results, one occasionally outperforms the other.

##### 1.1.3 Simulation Scheme 2

We now examine simulation scheme 2, which introduces correlations between features. Recall that in simulation scheme 1, *GenF* and *sAFT* performed similarly and better than the unadjusted approach, and *ξ* outperformed *ω*. In this section, we demonstrate that the same conclusions can be made for scheme 2. However, although the results have a similar trend, the AUC and Youden values for scheme 2 are lower than those in scheme 1 likely due to the complexity in the data introduced by the correlation structure between features. The AUCs for scheme 2 using *ξ* are shown in SI Table 9. Similar to scheme 1, we observe that our adjusted methods have a clear advantage over the unadjusted method and this difference becomes greater as the censoring fraction increases. We observe that *GenF* results in higher AUCs for LN and LL2, which differs slightly from scheme 1 results, but *GenF* and *sAFT* perform very similarly in the remaining schemes. The AUCs using *ω* were observed to be much lower than those obtained using *ξ* in each case (data not shown); in fact, the AUCs for *ω* ranged from 0.50 to 0.57 which suggests that its selection capability is only slightly better than a coin flip. Thus, *ξ* is seen to be optimal approach for feature selection. Next, we examine the Youden index displayed in SI Table 10. We note that the Youden indices based on *ω* are lower than those based on *ξ* in every single case, an observation similar to that in scheme 1. In fact, in many cases, the Youden index for *ω* is 0. We also observe from these results that *GenF* and *sAFT* outperform the unadjusted method in practically every scenario and particularly for higher censoring. *GenF* and *sAFT* perform similarly in many cases, but as indicated by the AUC results, one occasionally outperforms the other. The simulation results from both schemes showed *GenF* either outperformed or matched the performance of LN, LL and W. Hence, we focus only on *GenF* in the examples in the remainder of this paper.

#### 1.2 Evaluating the prediction algorithm

Simulated data was generated using the approach outlined in §6.1 for schemes 1 and 2 based on LN, LL1, LL2, W1 and W2 models for 33%, 67% and 80% censoring. Each data set contained 200 observations and 5,000 mock features and model parameters were chosen appropriately to result in survival times that mimicked actual survival times (say in months or years). Training and test sets were obtained using a 2:1 split. For each combination of simulation scheme, model and censoring proportion, a total of 25 different random splits were generated and the predictive performance of ACPR-AFT (using *α* = 0.5) was evaluated on each data set using *GenF* or *sAFT* and compared with the unadjusted method, CPR-AFT. Median summaries for different measures of prediction accuracy are presented in SI Tables 10-15 where *MSE_T R_* and *MSE_T E_* refer to the MSE for the training and test set, respectively. It is not surprising to note that the overall performance of training sets is better than that of test sets across all parameters considered for both CPR-AFT and ACPR-AFT. However, as the censoring fraction increases, a significant improvement is noted in the predictive performance (AUC and MSE) of ACPR-AFT (both *GenF* and *sAFT*) compared to CPR-AFT for the test sets. This improved performance is observed under both simulation schemes for each of the five different models under consideration. These results, thus, highlight the superiority of ACPR-AFT under a variety of data generating mechanisms encountered in practice.

**Table 1:** Simulation Scheme 1

| Censoring | Method | $\omega$ | | $\xi$ | |
| --- | --- | --- | --- | --- | --- |
|  |  | Youden | AUC | Youden | AUC |
| 33 % | ACPR-AFT LN | .19 | .67 | .65 | .94 |
|  | ACPR-AFT W | .21 | .68 | .65 | .94 |
|  | ACPR-AFT LL | .20 | .67 | .65 | .94 |
|  | ACPR-AFT *GenF* | .33 | .75 | .74 | .96 |
|  | CPR-AFT | .12 | .64 | .48 | .88 |
| 80% | ACPR-AFT LN | .16 | .63 | .30 | .74 |
|  | ACPR-AFT W | .16 | .63 | .30 | .74 |
|  | ACPR-AFT LL | .16 | .63 | .30 | .74 |
|  | ACPR-AFT *GenF* | .44 | .80 | .49 | .84 |
|  | CPR-AFT | .03 | .51 | .09 | .62 |

**Table 2:** Simulation Scheme 1: AUC (*ξ*)

| Censoring | Method | LN | LL1 | LL2 | W1 | W2 |
| --- | --- | --- | --- | --- | --- | --- |
| 33 % | ACPR-AFT *GenF* | .91 | .92 | .69 | .95 | .89 |
|  | ACPR-AFT *sAFT* | .91 | .92 | .76 | .97 | .92 |
|  | CPR-AFT | .88 | .89 | .74 | .93 | .89 |
| 67 % | ACPR-AFT *GenF* | .87 | .89 | .72 | .90 | .76 |
|  | ACPR-AFT *sAFT* | .86 | .84 | .71 | .95 | .84 |
|  | CPR-AFT | .80 | .78 | .65 | .85 | .72 |
| 80 % | ACPR-AFT *GenF* | .85 | .84 | .75 | .87 | .64 |
|  | ACPR-AFT *sAFT* | .81 | .77 | .68 | .92 | .70 |
|  | CPR-AFT | .72 | .66 | .59 | .77 | .56 |

**Table 3:** Simulation Scheme 1: AUC (*ω*)

| Censoring | Method | LN | LL1 | LL2 | W1 | W2 |
| --- | --- | --- | --- | --- | --- | --- |
| 33 % | ACPR-AFT *GenF* | .67 | .64 | .52 | .66 | .56 |
|  | ACPR-AFT *sAFT* | .68 | .64 | .55 | .71 | .54 |
|  | CPR-AFT | .60 | .57 | .53 | .62 | .54 |
| 67 % | ACPR-AFT *GenF* | .71 | .63 | .57 | .62 | .52 |
|  | ACPR-AFT *sAFT* | .66 | .55 | .56 | .73 | .55 |
|  | CPR-AFT | .54 | .51 | .53 | .54 | .50 |
| 80 % | ACPR-AFT *GenF* | .68 | .63 | .60 | .62 | .51 |
|  | ACPR-AFT *sAFT* | .59 | .54 | .54 | .75 | .53 |
|  | CPR-AFT | .51 | .51 | .51 | .53 | .50 |

**Table 4:** Simulation Scheme 1: Youden Index

| Model | Method | 33% cens. |  | 67% cens. |  | 80% cens. |  |
| --- | --- | --- | --- | --- | --- | --- | --- |
| | | $\omega$ | $\xi$ | $\omega$ | $\xi$ | $\omega$ | $\xi$ |
| LN | ACPR-AFT *GenF* | .21 | .52 | .29 | .55 | .22 | .52 |
|  | ACPR-AFT *sAFT* | .21 | .52 | .19 | .52 | .10 | .48 |
|  | CPR-AFT | .10 | .50 | .03 | .41 | .01 | .17 |
| LL1 | ACPR-AFT *GenF* | .15 | .56 | .15 | .54 | .14 | .45 |
|  | ACPR-AFT *sAFT* | .15 | .56 | .05 | .43 | .04 | .24 |
|  | CPR-AFT | .06 | .50 | .01 | .21 | .01 | .07 |
| LL2 | ACPR-AFT *GenF* | .02 | .20 | .07 | .23 | .11 | .31 |
|  | ACPR-AFT *sAFT* | .05 | .29 | .06 | .20 | .04 | .14 |
|  | CPR-AFT | .03 | .23 | .03 | .10 | .01 | .05 |
| W1 | ACPR-AFT *GenF* | .18 | .62 | .12 | .52 | .14 | .50 |
|  | ACPR-AFT *sAFT* | .24 | .70 | .31 | .65 | .34 | .61 |
|  | CPR-AFT | .11 | .56 | .04 | .40 | .03 | .21 |
| W2 | ACPR-AFT *GenF* | .04 | .45 | .02 | .22 | .01 | .10 |
|  | ACPR-AFT *sAFT* | .07 | .52 | .05 | .39 | .04 | .18 |
|  | CPR-AFT | .03 | .43 | .01 | .12 | .01 | .02 |

**Table 5:**
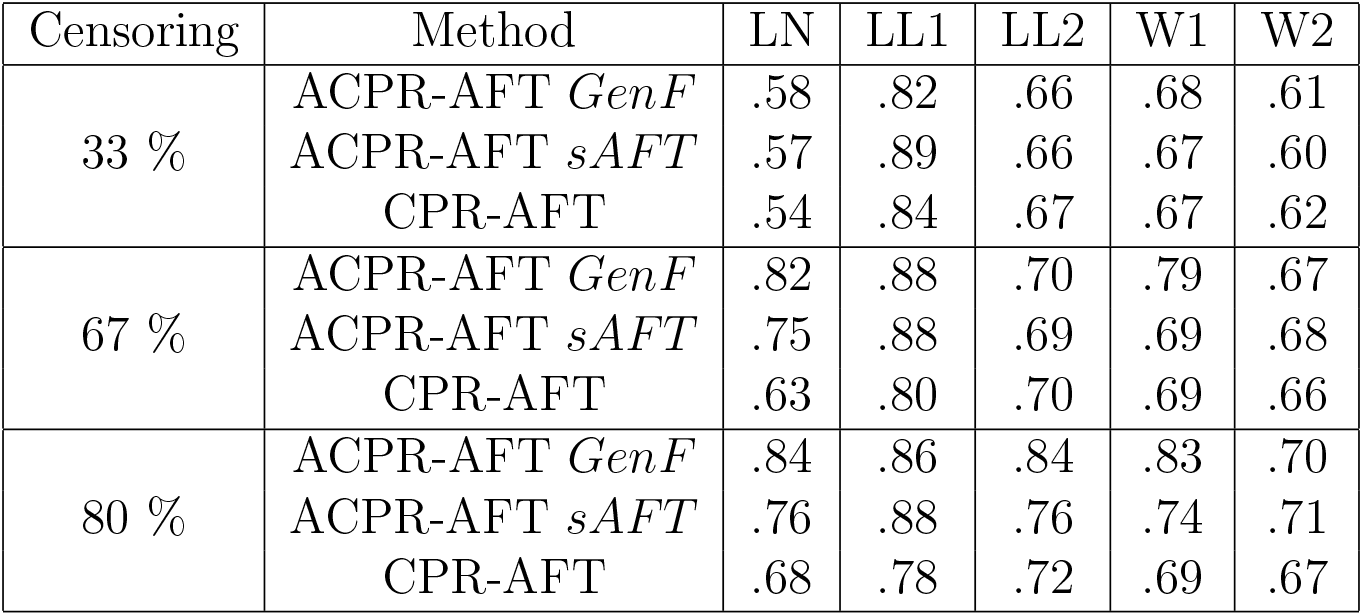
Simulation Scheme 2: AUC (*ξ*)

**Table 6:** Simulation Scheme 2: Youden Index

| Model | Method | 33% cens. |  | 67% cens. |  | 80% cens. |  |
| --- | --- | --- | --- | --- | --- | --- | --- |
| | | $\omega$ | $\xi$ | $\omega$ | $\xi$ | $\omega$ | $\xi$ |
| LN | ACPR-AFT *GenF* | 0 | .01 | 0 | .11 | 0 | .13 |
|  | ACPR-AFT *sAFT* | .01 | .01 | .01 | .09 | 0 | .09 |
|  | CPR-AFT | 0 | .01 | 0 | .01 | 0 | .03 |
| LL1 | ACPR-AFT *GenF* | .05 | .19 | .06 | .24 | .06 | .20 |
|  | ACPR-AFT *sAFT* | .06 | .24 | .04 | .23 | .04 | .22 |
|  | CPR-AFT | .02 | .16 | .01 | .10 | 0 | .08 |
| LL2 | ACPR-AFT *GenF* | 0 | .03 | 0 | .04 | 0 | .13 |
|  | ACPR-AFT *sAFT* | 0 | .03 | 0 | .03 | 0 | .06 |
|  | CPR-AFT | 0 | .03 | 0 | .03 | 0 | .05 |
| W1 | ACPR-AFT *GenF* | .01 | .04 | .01 | .09 | .02 | .15 |
|  | ACPR-AFT *sAFT* | .01 | .03 | .02 | .05 | .01 | .07 |
|  | CPR-AFT | 0 | .03 | .01 | .04 | 0 | .04 |
| W2 | ACPR-AFT *GenF* | .01 | .02 | 0 | .03 | 0 | .04 |
|  | ACPR-AFT *sAFT* | .01 | .02 | 0 | .04 | 0 | .03 |
|  | CPR-AFT | 0 | .02 | 0 | .02 | 0 | .02 |

**Table 7:** Simulation Scheme 1: 33% Censoring

| Model | Measure | ACPR-AFT<br>*GenF* | ACPR-AFT<br>*sAFT* | CPR-AFT |
| --- | --- | --- | --- | --- |
| LN, Scheme 1 | $R_{COMP}^2$ | 100 | 99.9 | 99.9 |
| | $MSE_{TR}$ | 0.007 | 0.009 | 0.008 |
| | AUC, $t = 2$ | 0.94 | 0.96 | 0.92 |
| | AUC, $t = 3$ | 0.91 | 0.93 | 0.89 |
| | $MSE_{TE}$ | 12.7 | 12.8 | 13.4 |
| | AUC, $t = 2$ | 0.94 | 0.92 | 0.91 |
| | AUC, $t = 3$ | 0.93 | 0.95 | 0.94 |
| LL1, Scheme 1 | $R_{COMP}^2$ | 100 | 100 | 100 |
| | $MSE_{TR}$ | 0.0005 | 0.0006 | 0.0006 |
| | AUC, $t = 2$ | 0.99 | 0.99 | 0.99 |
| | AUC, $t = 3$ | 0.99 | 0.99 | 0.99 |
| | $MSE_{TE}$ | 2.05 | 2.05 | 2.06 |
| | AUC, $t = 2$ | 0.97 | 0.97 | 0.97 |
| | AUC, $t = 3$ | 0.97 | 0.96 | 0.96 |
| LL2, Scheme 1 | $R_{COMP}^2$ | 99.8 | 100 | 99.8 |
| | $MSE_{TR}$ | 0.03 | 0.001 | 0.03 |
| | AUC, $t = 2$ | 0.98 | 0.99 | 0.97 |
| | AUC, $t = 3$ | 0.98 | 0.98 | 0.97 |
| | $MSE_{TE}$ | 2.93 | 2.95 | 3.28 |
| | AUC, $t = 2$ | 0.88 | 0.88 | 0.88 |
| | AUC, $t = 3$ | 0.93 | 0.95 | 0.94 |
| W1, Scheme 1 | $R_{COMP}^2$ | 100 | 100 | 99.5 |
| | $MSE_{TR}$ | 0.001 | 0.001 | 0.001 |
| | AUC, $t = 2$ | 0.98 | 0.98 | 0.99 |
| | AUC, $t = 3$ | 0.98 | 0.99 | 0.98 |
| | $MSE_{TE}$ | 4.21 | 4.14 | 4.94 |
| | AUC, $t = 2$ | 0.99 | 0.99 | 0.98 |
| | AUC, $t = 3$ | 0.98 | 0.98 | 0.98 |
| W2, Scheme 1 | $R_{COMP}^2$ | 100 | 100 | 98.9 |
| | $MSE_{TR}$ | 0.001 | 0.001 | 0.05 |
| | AUC, $t = 2$ | 0.98 | 0.98 | 0.98 |
| | AUC, $t = 3$ | 0.97 | 0.98 | 0.96 |
| | $MSE_{TE}$ | 7.52 | 7.66 | 8.68 |
| | AUC, $t = 2$ | 0.86 | 0.87 | 0.86 |
| | AUC, $t = 3$ | 0.92 | 0.94 | 0.92 |

**Table 8:** Simulation Scheme 2: 33% Censoring

| Model | Measure | ACPR-AFT<br>*GenF* | ACPR-AFT<br>*sAFT* | CPR-AFT |
| --- | --- | --- | --- | --- |
| LN, Scheme 2 | $R^2_{COMP}$ | 100 | 98.2 | 98.1 |
| | $MSE_{TR}$ | 0.006 | 0.179 | 0.228 |
| | AUC, $t = 2$ | 0.75 | 0.76 | 0.76 |
| | AUC, $t = 3$ | 0.89 | 0.89 | 0.9 |
| | $MSE_{TE}$ | 9.3 | 9.3 | 9.2 |
| | AUC, $t = 2$ | 0.89 | 0.91 | 0.91 |
| | AUC, $t = 3$ | 0.88 | 0.9 | 0.84 |
| LL1, Scheme 2 | $R^2_{COMP}$ | 99.9 | 99.9 | 100 |
| | $MSE_{TR}$ | 0.002 | 0.002 | 0.001 |
| | AUC, $t = 2$ | 0.71 | 0.7 | 0.66 |
| | AUC, $t = 3$ | 0.71 | 0.71 | 0.67 |
| | $MSE_{TE}$ | 3.12 | 3.11 | 3.14 |
| | AUC, $t = 2$ | 0.99 | 0.98 | 0.97 |
| | AUC, $t = 3$ | 0.97 | 0.97 | 0.96 |
| LL2, Scheme 2 | $R^2_{COMP}$ | 97.7 | 98 | 97.6 |
| | $MSE_{TR}$ | 0.056 | 0.097 | 0.092 |
| | AUC, $t = 2$ | 0.56 | 0.56 | 0.56 |
| | AUC, $t = 3$ | 0.59 | 0.58 | 0.57 |
| | $MSE_{TE}$ | 2.3 | 2.3 | 2.3 |
| | AUC, $t = 2$ | 0.98 | 0.97 | 0.95 |
| | AUC, $t = 3$ | 0.86 | 0.96 | 0.89 |
| W1, Scheme 2 | $R^2_{COMP}$ | 98.2 | 98 | 98.1 |
| | $MSE_{TR}$ | 0.08 | 0.1 | 0.09 |
| | AUC, $t = 2$ | 0.84 | 0.84 | 0.85 |
| | AUC, $t = 3$ | 0.79 | 0.8 | 0.8 |
| | $MSE_{TE}$ | 4.36 | 4.04 | 4.02 |
| | AUC, $t = 2$ | 0.98 | 0.99 | 0.99 |
| | AUC, $t = 3$ | 0.95 | 0.98 | 0.98 |
| W2, Scheme 2 | $R^2_{COMP}$ | 99.9 | 98 | 98 |
| | $MSE_{TR}$ | 0.005 | 0.127 | 0.114 |
| | AUC, $t = 2$ | 0.91 | 0.91 | 0.92 |
| | AUC, $t = 3$ | 0.87 | 0.88 | 0.88 |
| | $MSE_{TE}$ | 6.12 | 5.29 | 5.3 |
| | AUC, $t = 2$ | 0.98 | 0.98 | 0.98 |
| | AUC, $t = 3$ | 0.98 | 0.98 | 0.97 |

**Table 9:** Simulation Scheme 1: 67% Censoring

| Model | Measure | ACPR-AFT<br>*GenF* | ACPR-AFT<br>*sAFT* | CPR-AFT |
| --- | --- | --- | --- | --- |
| LN, Scheme 1 | $R^2_{COMP}$ | 100 | 100 | 99.9 |
| | $MSE_{TR}$ | 0.003 | 0.006 | 0.007 |
| | AUC, $t = 2$ | 0.82 | 0.88 | 0.81 |
| | AUC, $t = 3$ | 0.82 | 0.86 | 0.79 |
| | $MSE_{TE}$ | 10.9 | 11.6 | 12.7 |
| | AUC, $t = 2$ | 0.86 | 0.84 | 0.84 |
| | AUC, $t = 3$ | 0.85 | 0.85 | 0.84 |
| LL1, Scheme 1 | $R^2_{COMP}$ | 100 | 100 | 100 |
| | $MSE_{TR}$ | 0.001 | 0.00001 | 0.003 |
| | AUC, $t = 2$ | 0.98 | 0.97 | 0.97 |
| | AUC, $t = 3$ | 0.96 | 0.96 | 0.94 |
| | $MSE_{TE}$ | 1.35 | 1.25 | 1.36 |
| | AUC, $t = 2$ | 0.94 | 0.86 | 0.84 |
| | AUC, $t = 3$ | 0.91 | 0.84 | 0.79 |
| LL2, Scheme 1 | $R^2_{COMP}$ | 100 | 100 | 100 |
| | $MSE_{TR}$ | 0.0002 | 0.0003 | 0.0004 |
| | AUC, $t = 2$ | 0.87 | 0.89 | 0.85 |
| | AUC, $t = 3$ | 0.93 | 0.94 | 0.91 |
| | $MSE_{TE}$ | 1.93 | 1.89 | 1.99 |
| | AUC, $t = 2$ | 0.88 | 0.86 | 0.84 |
| | AUC, $t = 3$ | 0.8 | 0.84 | 0.79 |
| W1, Scheme 1 | $R^2_{COMP}$ | 100 | 100 | 99.5 |
| | $MSE_{TR}$ | 0.003 | 0.0003 | 0.01 |
| | AUC, $t = 2$ | 0.97 | 0.97 | 0.96 |
| | AUC, $t = 3$ | 0.93 | 0.93 | 0.92 |
| | $MSE_{TE}$ | 3.7 | 3.79 | 4.09 |
| | AUC, $t = 2$ | 0.92 | 0.9 | 0.86 |
| | AUC, $t = 3$ | 0.92 | 0.88 | 0.74 |
| W2, Scheme 1 | $R^2_{COMP}$ | 100 | 100 | 100 |
| | $MSE_{TR}$ | 0.0004 | 0.00001 | 0.0005 |
| | AUC, $t = 2$ | 0.96 | 0.97 | 0.96 |
| | AUC, $t = 3$ | 0.98 | 0.97 | 0.97 |
| | $MSE_{TE}$ | 5.52 | 5.51 | 5.57 |
| | AUC, $t = 2$ | 0.89 | 0.89 | 0.86 |
| | AUC, $t = 3$ | 0.9 | 0.85 | 0.79 |

**Table 10:** Simulation Scheme 2: 67% Censoring

| Model | Measure | ACPR-AFT<br>*GenF* | ACPR-AFT<br>*sAFT* | CPR-AFT |
| --- | --- | --- | --- | --- |
| LN, Scheme 2 | $R^2_{COMP}$ | 100 | 100 | 100 |
| | $MSE_{TR}$ | 0.000002 | 0.000002 | 0.000002 |
| | AUC, $t = 2$ | 0.8 | 0.79 | 0.79 |
| | AUC, $t = 3$ | 0.92 | 0.92 | 0.92 |
| | $MSE_{TE}$ | 4.9 | 4.9 | 5 |
| | AUC, $t = 2$ | 0.85 | 0.87 | 0.76 |
| | AUC, $t = 3$ | 0.87 | 0.87 | 0.83 |
| LL1, Scheme 2 | $R^2_{COMP}$ | 100 | 100 | 100 |
| | $MSE_{TR}$ | 0.00003 | 0.00004 | 0.0009 |
| | AUC, $t = 2$ | 0.94 | 0.9 | 0.79 |
| | AUC, $t = 3$ | 0.79 | 0.8 | 0.74 |
| | $MSE_{TE}$ | 1.12 | 1.11 | 1.14 |
| | AUC, $t = 2$ | 0.95 | 0.9 | 0.89 |
| | AUC, $t = 3$ | 0.87 | 0.84 | 0.8 |
| LL2, Scheme 2 | $R^2_{COMP}$ | 97.5 | 97.3 | 97.5 |
| | $MSE_{TR}$ | 0.016 | 0.046 | 0.061 |
| | AUC, $t = 2$ | 0.7 | 0.7 | 0.56 |
| | AUC, $t = 3$ | 0.72 | 0.7 | 0.58 |
| | $MSE_{TE}$ | 1.45 | 1.49 | 1.52 |
| | AUC, $t = 2$ | 0.95 | 0.86 | 0.76 |
| | AUC, $t = 3$ | 0.73 | 0.87 | 0.78 |
| W1, Scheme 2 | $R^2_{COMP}$ | 100 | 100 | 100 |
| | $MSE_{TR}$ | 0.001 | 0.001 | 0.001 |
| | AUC, $t = 2$ | 0.98 | 0.98 | 0.97 |
| | AUC, $t = 3$ | 0.96 | 0.96 | 0.95 |
| | $MSE_{TE}$ | 3.88 | 3.96 | 3.97 |
| | AUC, $t = 2$ | 0.94 | 0.92 | 0.9 |
| | AUC, $t = 3$ | 0.9 | 0.89 | 0.83 |
| W2, Scheme 2 | $R^2_{COMP}$ | 97.7 | 97.5 | 97.5 |
| | $MSE_{TR}$ | 0.05 | 0.06 | 0.08 |
| | AUC, $t = 2$ | 0.92 | 0.91 | 0.86 |
| | AUC, $t = 3$ | 0.95 | 0.93 | 0.89 |
| | $MSE_{TE}$ | 4.98 | 5 | 5 |
| | AUC, $t = 2$ | 0.98 | 0.93 | 0.93 |
| | AUC, $t = 3$ | 0.92 | 0.91 | 0.9 |

**Table 11:**
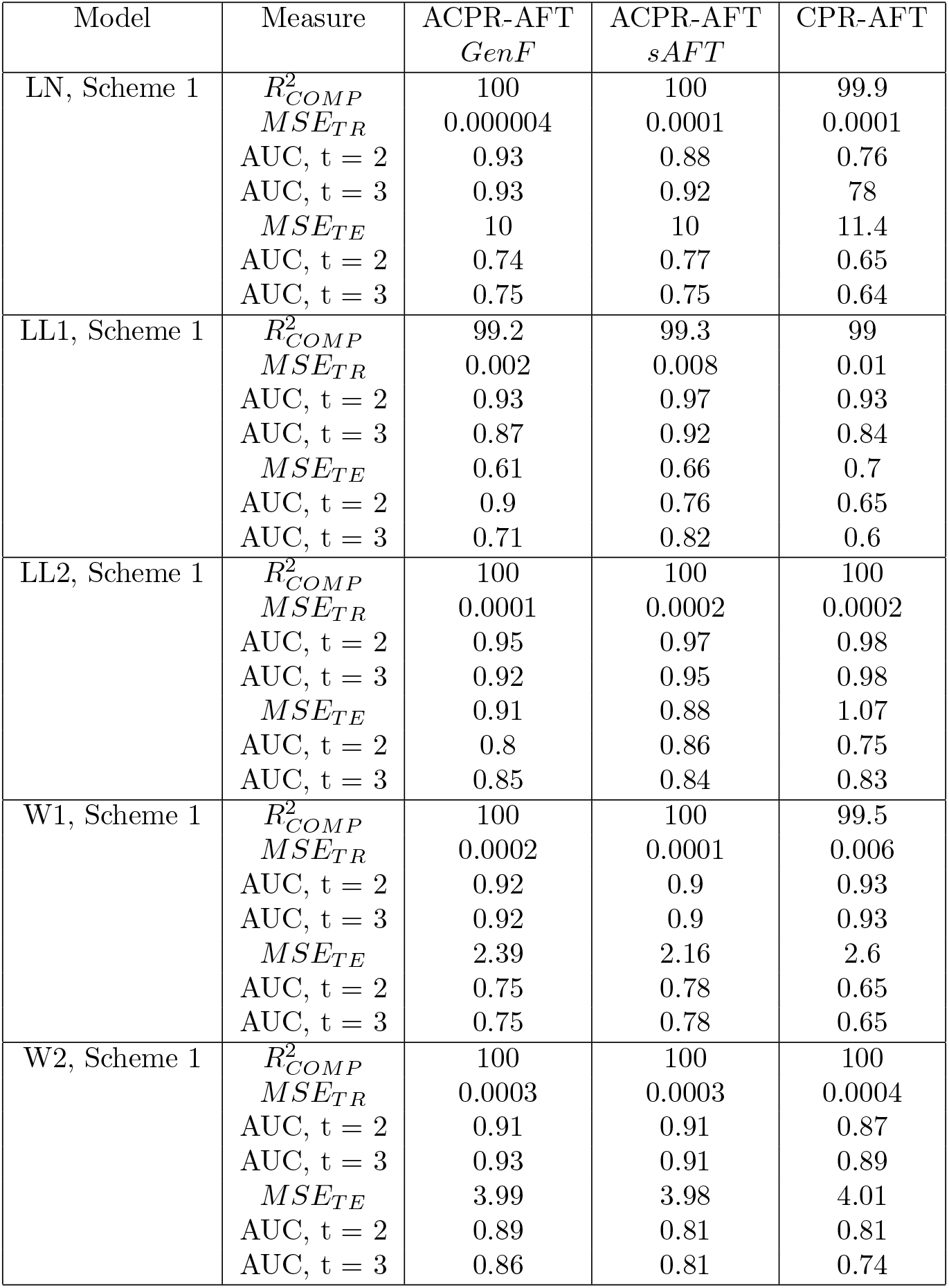
Simulation Scheme 1: 80% Censoring

**Table 12:** Simulation Scheme 2: 80% Censoring

| Model | Measure | ACPR-AFT<br>*GenF* | ACPR-AFT<br>*sAFT* | CPR-AFT |
| --- | --- | --- | --- | --- |
| LN, Scheme 2 | $R_{COMP}^2$ | 98.9 | 98.4 | 98.1 |
| | $MSE_{TR}$ | 0.035 | 0.12 | 0.14 |
| | AUC, $t = 2$ | 0.73 | 0.74 | 0.73 |
| | AUC, $t = 3$ | 0.83 | 0.82 | 0.79 |
| | $MSE_{TE}$ | 3.76 | 3.59 | 3.76 |
| | AUC, $t = 2$ | 0.89 | 0.95 | 0.84 |
| | AUC, $t = 3$ | 0.95 | 0.89 | 0.86 |
| LL1, Scheme 2 | $R_{COMP}^2$ | 100 | 100 | 100 |
| | $MSE_{TR}$ | 0.0002 | 0.0002 | 0.0005 |
| | AUC, $t = 2$ | 0.95 | 0.97 | 0.93 |
| | AUC, $t = 3$ | 0.94 | 0.96 | 0.92 |
| | $MSE_{TE}$ | 1.72 | 1.72 | 1.75 |
| | AUC, $t = 2$ | 0.68 | 0.79 | 0.66 |
| | AUC, $t = 3$ | 0.7 | 0.83 | 0.55 |
| LL2, Scheme 2 | $R_{COMP}^2$ | 100 | 100 | 97.7 |
| | $MSE_{TR}$ | 0.0003 | 0.00001 | 0.026 |
| | AUC, $t = 2$ | 0.86 | 0.89 | 0.68 |
| | AUC, $t = 3$ | 0.81 | 0.9 | 0.63 |
| | $MSE_{TE}$ | 1.23 | 1.13 | 1.33 |
| | AUC, $t = 2$ | 0.71 | 0.84 | 0.61 |
| | AUC, $t = 3$ | 0.62 | 0.91 | 0.61 |
| W1, Scheme 2 | $R_{COMP}^2$ | 100 | 100 | 100 |
| | $MSE_{TR}$ | 0.00002 | 0.00003 | 0.00002 |
| | AUC, $t = 2$ | 0.96 | 0.95 | 0.93 |
| | AUC, $t = 3$ | 0.96 | 0.95 | 0.93 |
| | $MSE_{TE}$ | 6.17 | 4.8 | 4.55 |
| | AUC, $t = 2$ | 0.75 | 0.88 | 0.68 |
| | AUC, $t = 3$ | 0.75 | 0.88 | 0.68 |
| W2, Scheme 2 | $R_{COMP}^2$ | 100 | 100 | 100 |
| | $MSE_{TR}$ | 0.0006 | 0.0007 | 0.0009 |
| | AUC, $t = 2$ | 0.98 | 0.99 | 0.98 |
| | AUC, $t = 3$ | 0.98 | 0.98 | 0.97 |
| | $MSE_{TE}$ | 4.73 | 4.29 | 5.25 |
| | AUC, $t = 2$ | 0.9 | 0.9 | 0.86 |
| | AUC, $t = 3$ | 0.79 | 0.82 | 0.66 |

**Figure 1:**
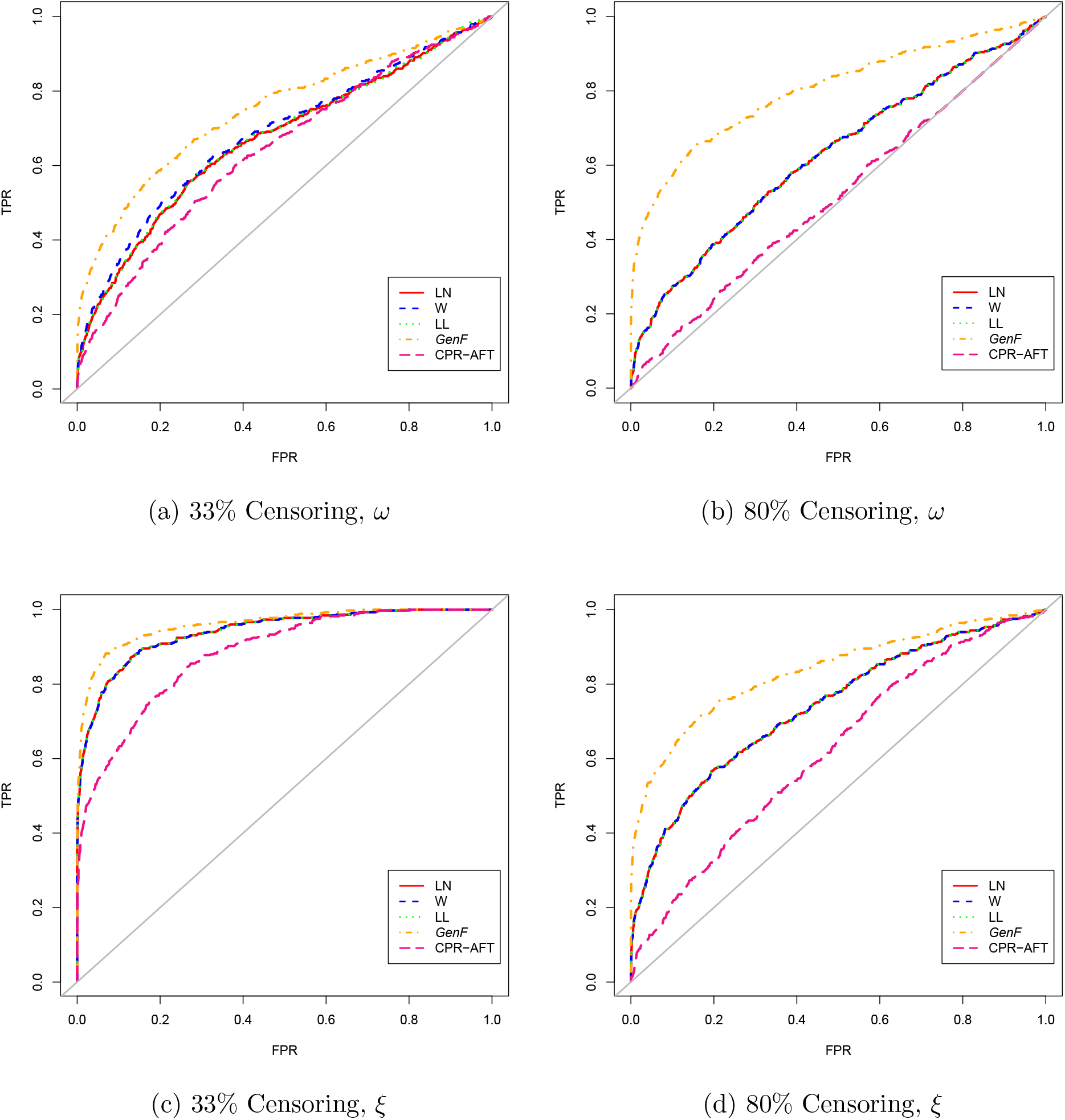
ROC Curves for Simulation Scheme 1, Comparison of *GenF* -based ACPR-AFT vs. its special cases and CPR-AFT for *ω* (a, b) and *ξ* (c, d).

**Figure 2:**
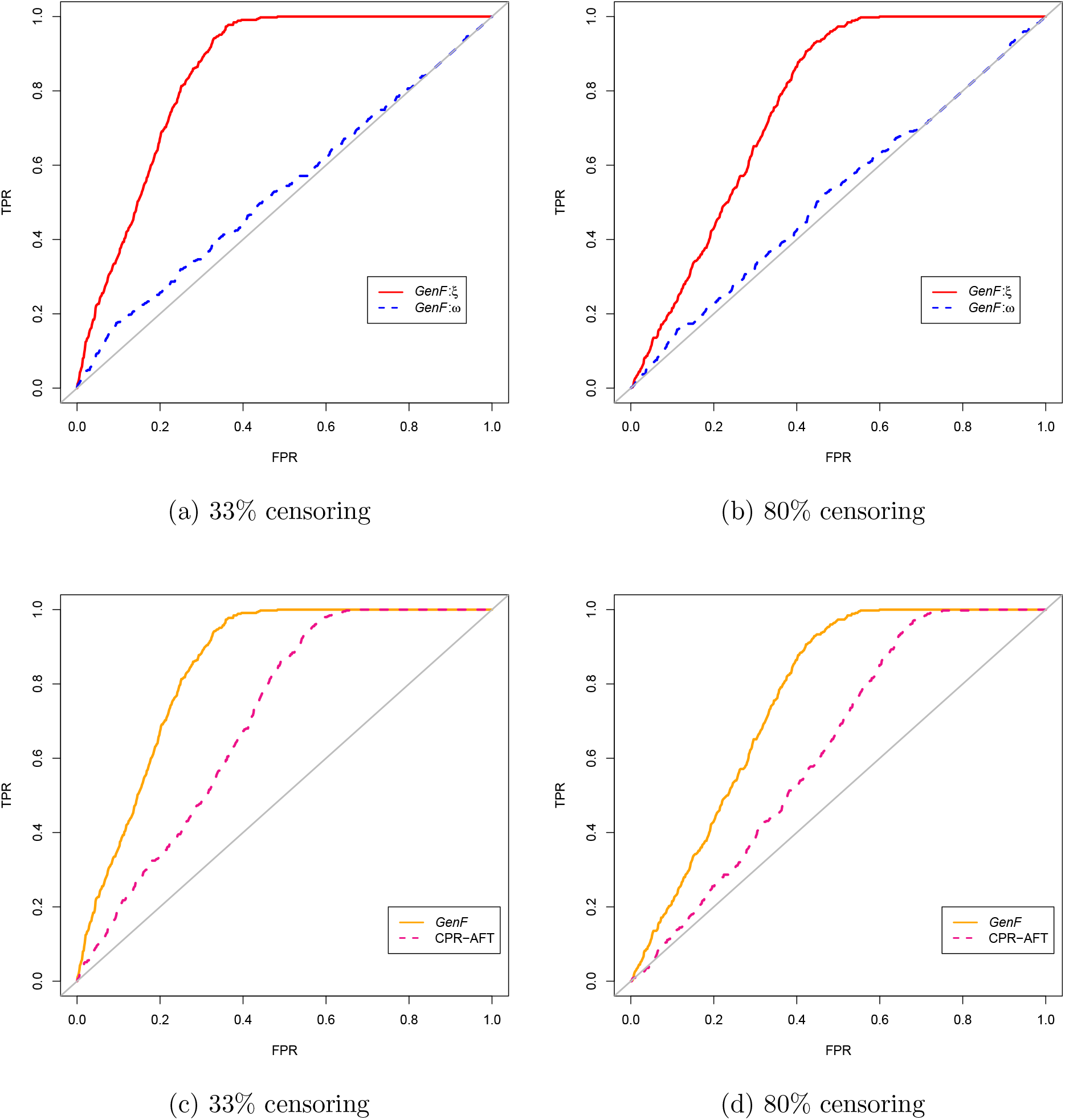
ROC Curves for Simulation Scheme 2, *ξ* vs. *ω* for *GenF* -based ACPR-AFT (a, b) and *GenF* -based ACPR-AFT vs. CPR-AFT using *ξ* (c, d)

### 2 Data sets

#### 2.1 Data sets

- Head & Neck squamous cell carcinoma (HNSCC): This data set was published by TCGA and contains the expression profiles of 19,341 genes obtained using RNA sequencing for 221 subjects with cancers of the oral cavity, a subgroup of HNSCC. Pre-processed gene expression data (RSEM values, Li & Dewey, 2011) was downloaded from the Broad Institute (http://gdac.broadinstitute.org) and further normalized using the log_2_(*x* + 1) transformation which accounts for exact zeros. A gene was included in the analyses only if (i) if 50% of patients have expression values for that gene, and (ii) protein expression of that gene was observed in at least one head and neck cancer sample in the Human Protein Atlas database (Uhlen et al., 2015). Overall survival is the endpoint of interest.
- Glioblastoma (GBM): This data set was published by TCGA and contains the methylation profiles (beta values) for 280 tumor samples obtained using the Infinium HumanMethylation27 platform. The beta values were normalized using the logit transformation. For genes with multiple methylation probes, the probe most negatively correlated with expression is used. Overall survival is the endpoint of interest.
- Ovarian cancer: Tothill et al. (2008) studied the relationship between gene expression and overall survival (OS) and recurrence-free survival (RFS) in ovarian cancer using tumor samples from 282 subjects and Affymetrix U133 Plus 2 microarrays. Affymetrix control probe sets and samples with missing survival data were removed from the RMA normalized data set (Irizarry et al., 2003). For OS, a coefficient of variation threshold of 35% was used to remove probe sets exhibiting low variation across tumor samples and resulted in 24,736 probe sets for 273 subjects. For RFS, a coefficient of variation threshold of 50% resulted in 13,696 probe sets for 276 subjects. Log_2_ transformed expression was used in all analyses.
- Oral cancer: Saintigny et al. (2011) studied 86 subjects enrolled in a clinical chemoprevention trial where the primary endpoint of interest was time to development of oral cancer. This RMA normalized and log_2_ transformed data set (Irizarry et al., 2003) contains the expression profiles of 12,776 probe sets obtained using the Human Gene 1.ST platform.

**Table 13:**
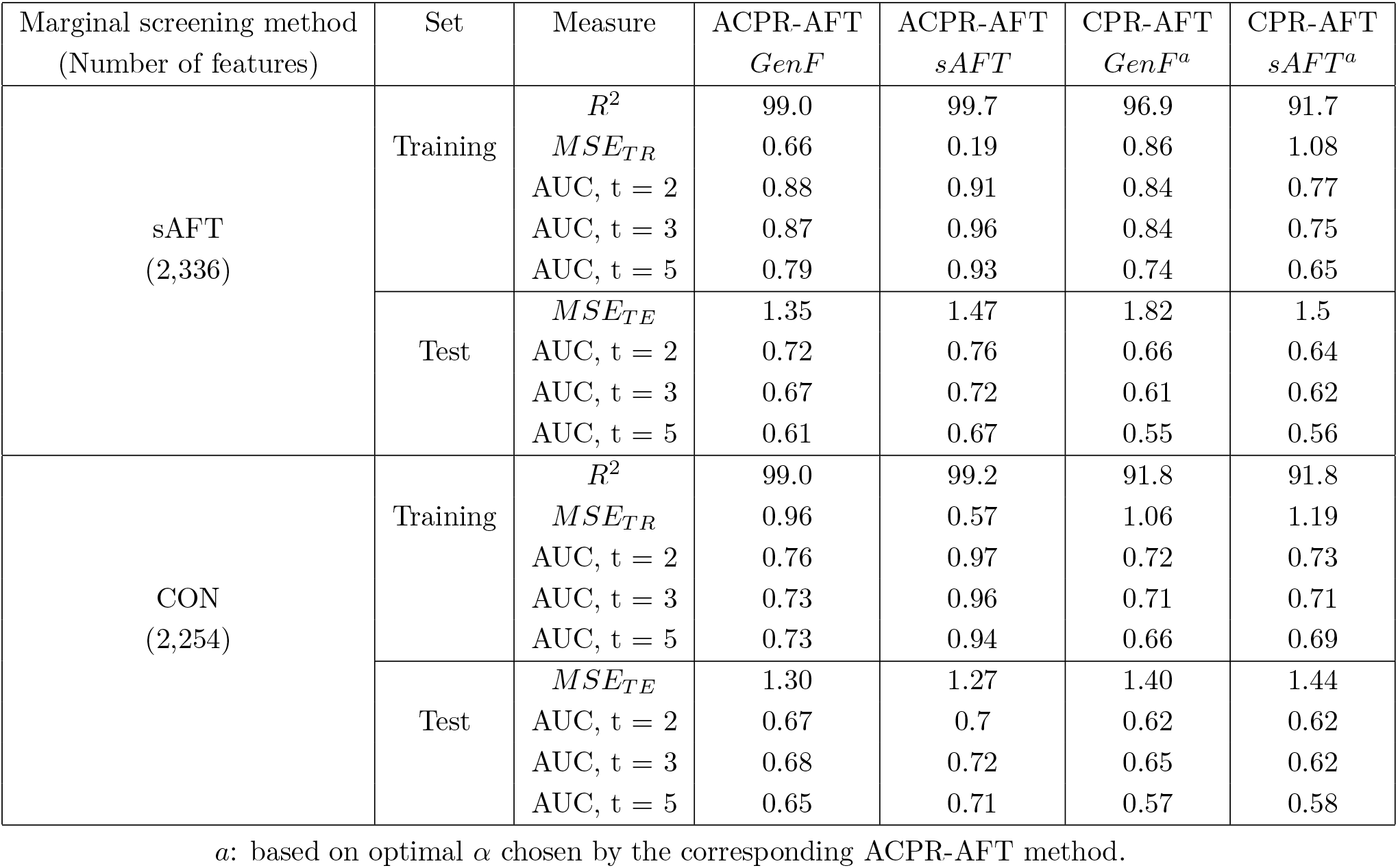
HNSCC Data: Summary of results on training and test sets

**Table 14:** GBM Data: Summary of results on training and test sets

| Marginal screening method<br>(Number of features) | Set | Measure | ACPR-AFT<br>*GenF* | ACPR-AFT<br>*sAFT* | CPR-AFT<br>*GenF*<sup>a</sup> | CPR-AFT<br>*sAFT*<sup>a</sup> |
| --- | --- | --- | --- | --- | --- | --- |
| sAFT<br>(3,605) | Training | $R^2$ | 87.9 | 90.3 | 85.0 | 85.6 |
| | | $MSE_{TR}$ | 1.10 | 0.89 | 0.90 | 0.90 |
|  |  | AUC, t = 2 | 0.63 | 0.74 | 0.63 | 0.71 |
|  |  | AUC, t = 3 | 0.70 | 0.83 | 0.70 | 0.69 |
|  |  | AUC, t = 5 | 0.95 | 0.94 | 0.80 | 0.86 |
| | Test | $MSE_{TE}$ | 1.42 | 1.33 | 1.45 | 1.35 |
|  |  | AUC, t = 2 | 0.61 | 0.62 | 0.61 | 0.61 |
|  |  | AUC, t = 3 | 0.69 | 0.66 | 0.68 | 0.67 |
|  |  | AUC, t = 5 | 0.82 | 0.82 | 0.82 | 0.77 |
| CON<br>(1,125) | Training | $R^2$ | 88.7 | 90.5 | 83.5 | 80.1 |
| | | $MSE_{TR}$ | 0.85 | 1.08 | 0.90 | 1.12 |
|  |  | AUC, t = 2 | 0.77 | 0.74 | 0.75 | 0.69 |
|  |  | AUC, t = 3 | 0.78 | 0.85 | 0.76 | 0.79 |
|  |  | AUC, t = 5 | 0.93 | 0.94 | 0.91 | 0.90 |
| | Test | $MSE_{TE}$ | 1.31 | 1.22 | 1.33 | 1.35 |
|  |  | AUC, t = 2 | 0.63 | 0.65 | 0.57 | 0.58 |
|  |  | AUC, t = 3 | 0.79 | 0.74 | 0.69 | 0.69 |
|  |  | AUC, t = 5 | 0.90 | 0.93 | 0.79 | 0.78 |

<sup>a</sup>: based on optimal $\alpha$ chosen by the corresponding ACPR-AFT method.

**Table 15:** Ovarian Cancer Data (OS): Summary of results on training and test sets

| Marginal screening method<br>(Number of features) | Set | Measure | ACPR-AFT<br>*GenF* | ACPR-AFT<br>*sAFT* | CPR-AFT<br>*GenF*<sup>a</sup> | CPR-AFT<br>*sAFT*<sup>a</sup> |
| --- | --- | --- | --- | --- | --- | --- |
| sAFT<br>(3,119) | Training | $R^2$ | 100 | 99.7 | 98.5 | 98.5 |
| | | $MSE_{TR}$ | 0.004 | 0.03 | 0.15 | 0.15 |
|  |  | AUC, t = 2 | 0.98 | .98 | 0.89 | 0.88 |
|  |  | AUC, t = 3 | 0.94 | .94 | 0.83 | 0.83 |
|  |  | AUC, t = 5 | 0.90 | .90 | 0.75 | 0.76 |
| | Test | $MSE_{TE}$ | 0.53 | 0.53 | 0.54 | 0.52 |
|  |  | AUC, t = 2 | 0.76 | 0.76 | 0.74 | 0.74 |
|  |  | AUC, t = 3 | 0.74 | 0.74 | 0.74 | 0.72 |
|  |  | AUC, t = 5 | 0.76 | 0.74 | 0.70 | 0.68 |
| CON<br>(2,485) | Training | $R^2$ | 99.9 | 99.0 | 98.7 | 98.7 |
| | | $MSE_{TR}$ | 0.14 | 0.14 | 0.17 | 0.17 |
|  |  | AUC, t = 2 | 0.95 | 0.95 | 0.88 | 0.88 |
|  |  | AUC, t = 3 | 0.93 | 0.92 | 0.82 | 0.82 |
|  |  | AUC, t = 5 | 0.90 | 0.90 | 0.77 | 0.77 |
| | Test | $MSE_{TE}$ | 0.66 | 0.66 | 0.68 | 0.68 |
|  |  | AUC, t = 2 | 0.74 | 0.74 | 0.70 | 0.70 |
|  |  | AUC, t = 3 | 0.71 | 0.72 | 0.68 | 0.68 |
|  |  | AUC, t = 5 | 0.81 | 0.82 | 0.71 | 0.73 |

<sup>a</sup>: based on optimal $\alpha$ chosen by the corresponding ACPR-AFT method.

**Table 16:** Ovarian Cancer Data (RFS): Summary of results on training and test sets

| Marginal screening method<br>(Number of features) | Set | Measure | ACPR-AFT<br>*GenF* | ACPR-AFT<br>*sAFT* | CPR-AFT<br>*GenF*<sup>a</sup> | CPR-AFT<br>*sAFT*<sup>a</sup> |
| --- | --- | --- | --- | --- | --- | --- |
| sAFT<br>(1,880) | Training | $R^2$ | 99.7 | 99.5 | 98.5 | 96.9 |
| | | $MSE_{TR}$ | 0.02 | 0.04 | 0.03 | 0.21 |
|  |  | AUC, t = 2 | 0.94 | 0.96 | 0.90 | 0.89 |
|  |  | AUC, t = 3 | 0.93 | 0.95 | 0.89 | 0.89 |
|  |  | AUC, t = 5 | 0.87 | 0.92 | 0.85 | 0.85 |
| | Test | $MSE_{TE}$ | 0.47 | 0.46 | 0.48 | 0.47 |
|  |  | AUC, t = 2 | 0.73 | 0.77 | 0.72 | 0.72 |
|  |  | AUC, t = 3 | 0.75 | 0.82 | 0.74 | 0.77 |
|  |  | AUC, t = 5 | 0.68 | 0.76 | 0.65 | 0.71 |
| CON<br>(1,682) | Training | $R^2$ | 99.9 | 99.7 | 98.9 | 96.5 |
| | | $MSE_{TR}$ | 0.007 | 0.21 | 0.007 | 0.22 |
|  |  | AUC, t = 2 | 0.94 | 0.91 | 0.92 | 0.85 |
|  |  | AUC, t = 3 | 0.93 | 0.94 | 0.89 | 0.88 |
|  |  | AUC, t = 5 | 0.89 | 0.92 | 0.85 | 0.85 |
| | Test | $MSE_{TE}$ | 0.41 | 0.43 | 0.50 | 0.49 |
|  |  | AUC, t = 2 | 0.74 | 0.76 | 0.71 | 0.72 |
|  |  | AUC, t = 3 | 0.80 | 0.82 | 0.76 | 0.75 |
|  |  | AUC, t = 5 | 0.78 | 0.79 | 0.74 | 0.80 |

<sup>a</sup>: based on optimal $\alpha$ chosen by the corresponding ACPR-AFT method.

**Table 17:** Oral Cancer Data: Summary of results on training and test sets

| Marginal screening method<br>(Number of features) | Set | Measure | ACPR-AFT<br>*GenF* | ACPR-AFT<br>*sAFT* | CPR-AFT<br>*GenF*<sup>a</sup> | CPR-AFT<br>*sAFT*<sup>a</sup> |
| --- | --- | --- | --- | --- | --- | --- |
| sAFT<br>(3,065) | Training | $R^2$ | 94.6 | 99 | 93.7 | 95.1 |
| | | $MSE_{TR}$ | 0.03 | 0.02 | 0.21 | 0.14 |
|  |  | AUC, t = 2 | 0.98 | 1.00 | 0.97 | 0.97 |
|  |  | AUC, t = 3 | 0.97 | 1.00 | 0.96 | 0.98 |
|  |  | AUC, t = 5 | 0.96 | 1.00 | 0.97 | 0.98 |
| | Test | $MSE_{TE}$ | 0.10 | 0.34 | 1.08 | 0.99 |
|  |  | AUC, t = 2 | 0.89 | 0.87 | 0.86 | 0.85 |
|  |  | AUC, t = 3 | 0.89 | 0.92 | 0.89 | 0.89 |
|  |  | AUC, t = 5 | 0.94 | 0.95 | 0.93 | 0.94 |
| CON<br>(4,652) | Training | $R^2$ | 95.6 | 95.6 | 89.3 | 88.3 |
| | | $MSE_{TR}$ | 0.01 | 0.12 | 0.40 | 0.4 |
|  |  | AUC, t = 2 | 0.96 | 0.97 | 0.95 | 0.94 |
|  |  | AUC, t = 3 | 0.98 | 0.99 | 0.97 | 0.97 |
|  |  | AUC, t = 5 | 0.97 | 0.99 | 0.96 | 0.98 |
| | Test | $MSE_{TE}$ | 0.36 | 0.44 | 1.20 | 1.17 |
|  |  | AUC, t = 2 | 0.81 | 0.83 | 0.82 | 0.82 |
|  |  | AUC, t = 3 | 0.86 | 0.88 | 0.86 | 0.86 |
|  |  | AUC, t = 5 | 0.90 | 0.92 | 0.90 | 0.91 |

<sup>a</sup>: based on optimal $\alpha$ chosen by the corresponding ACPR-AFT method.

**Figure 3:**
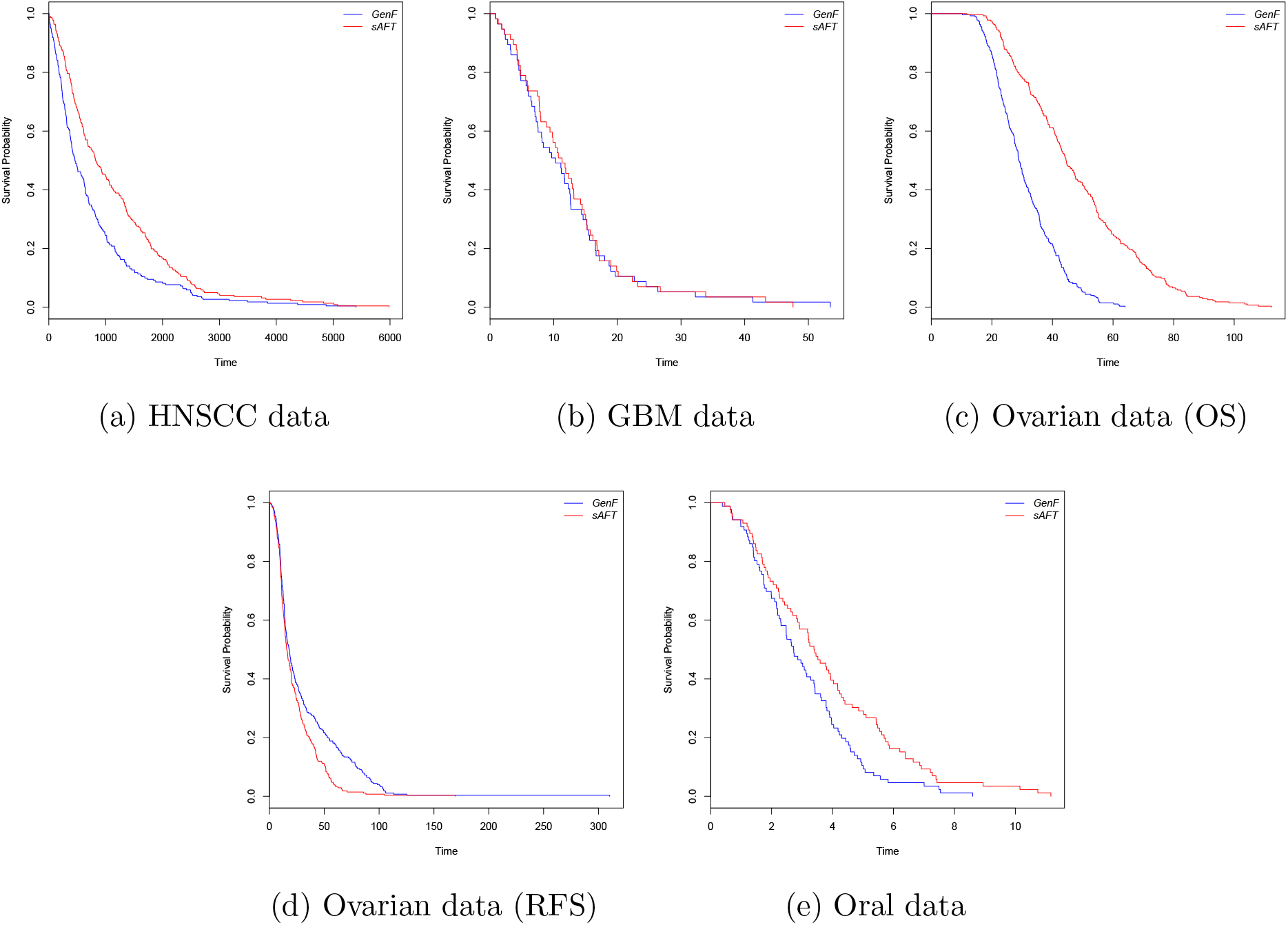
Predicted survival curves based on *PI*, *GenF* vs. *sAFT* -based ACPR-AFT

